# A Ratiometric Catalog of Protein Isoform Shifts in the Cardiac Fetal Gene Program

**DOI:** 10.1101/2024.04.09.588716

**Authors:** Yu Han, Shaonil Binti, Sara A. Wennersten, Boomathi Pandi, Dominic C. M. Ng, Edward Lau, Maggie P. Y. Lam

## Abstract

Pathological cardiac remodeling is associated with the reactivation of fetal genes, yet the extent of the heart’s fetal gene program and its impact on proteome compositions remain incompletely understood. Here, using a new proteome-wide protein ratio quantification strategy with mass spectrometry, we identify pervasive isoform usage shifts in fetal and postnatal mouse hearts, involving 145 pairs of highly homologous paralogs and alternative splicing-derived isoform proteins. Proteome-wide ratio comparisons readily rediscover hallmark fetal gene signatures in muscle contraction and glucose metabolism pathways, while revealing novel isoform usage in mitochondrial and gene expression proteins, including PPA1/PPA2, ANT1/ANT2, and PCBP1/PCBP2 switches. Paralogs with differential fetal usage tend to be evolutionarily recent, consistent with functional diversification. Alternative splicing adds another rich source of fetal isoform usage differences, involving PKM M1/M2, GLS-1 KGA/GAC, PDLIM5 long/short, and other spliceoforms. When comparing absolute protein proportions, we observe a partial reversion toward fetal gene usage in pathological hearts. In summary, we present a ratiometric catalog of paralogs and spliceoform pairs in the cardiac fetal gene program. More generally, the results demonstrate the potential of applying the proteome-wide ratio test concept to discover new regulatory modalities beyond differential gene expression.

## Introduction

The proteome undergoes tremendous transformations during development as it adapts to each organ’s changing energetic and physiologic needs. In the heart, a sudden shift occurs around birth. The fetal heart switches from mainly oxidizing carbohydrates as fuel in a relatively hypoxic environment, toward preferring fatty acids under high arterial oxygen levels in the postnatal niche. This is paralleled by other development and maturation milestones, such as increased contractile forces, decreased resilience to hypoxia, and the postnatal loss of regenerative capacity (Dimasi et al., 2023; Galdos et al., 2017). At the molecular level, these seismic changes are underpinned by a switch in the usage of multiple fetal variants of metabolic and contractile genes towards their postnatal counterparts (Franco et al., 1998; Hunkeler et al., 1991; Keller et al., 1995; Kuwahara et al., 2012; Lyons et al., 1990; Talman et al., 2018). Under pathological remodeling in hypertrophic and failing adult hearts, a remarkable reversal back to fetal gene expression has been widely observed (Cox and Marsh, 2014; Dirkx et al., 2013; Lowes et al., 2002; Oka et al., 2007; Rajabi et al., 2007; Razeghi et al., 2001). While it remains debated whether fetal gene reactivation is primarily adaptive (Rajabi et al., 2007; Taegtmeyer et al., 2010) or maladaptive (Dirkx et al., 2013) in nature, the fetal gene program has received broad interest for understanding key processes in cardiac development and disease. Many studies have successfully harnessed large-scale transcriptomics technologies to identify the trajectory of gene expression among thousands of cardiac genes and across multiple stages of fetal development (D’Antonio et al., 2022; DeLaughter et al., 2016). However, the trajectory of RNA levels provides only a modest prediction of protein abundance (Gygi et al., 1999; Payne, 2015; Srivastava et al., 2022; Yang et al., 2020) and cannot survey the effects of post-transcriptional and post-translational regulations on gene expression. Therefore, discovering the extent of cardiac fetal genes at the protein level is an important goal in the study of development and disease (Edwards et al., 2023; Gu et al., 2022).

Beyond the expression of individual genes, the fetal gene program of the heart has often been investigated by comparing the relative levels within pairs of proteins. An example is the reciprocal intensity of ɑ myosin heavy chain vs. β myosin heavy chain (Franco et al., 1998; Reiser et al., 2001), which bind to other proteins to form the sarcomere structure in different chambers of the heart. Other classic examples such as actin, BCL-2, and estrogen receptors have also shown that the ratio between two gene products can serve as the critical determinant of their biological function rather than the individual expression of either gene (Bergeron et al., 2010; Gelens et al., 2018; Warren et al., 2019; Zannoni et al., 2013). Mechanistically, this relationship can occur if two proteins participate in the formation of heteromultimeric complexes under stoichiometric constraints, or engage in mutually repressive feedback loops, compete for the binding of the same pool of substrates, or recognize identical genetic elements (Hart et al., 1999; Krikun et al., 2000; Li et al., 2014; Smink et al., 2009). In the example of the apoptotic regulators BAX and BCL-2, these two proteins form a heterodimer that together modulates the function of BAX. Rather than being regulated by the individual abundance of either protein, cells become sensitized to apoptosis when BAX/BCL-2 ratio is high, whereas resistance is conferred when BAX/BCL-2 ratio is low. In another example, the ratio of the Sp family transcription factors Sp1 and Sp3 helps regulate many biological processes in part because they target identical promoters (Apt et al., 1996; Hasegawa and Struhl, 2021). Critically, a recent large-scale analysis of genomics data has identified thousands of protein ratio quantitative trait loci (rQTL), i.e., polymorphisms that regulate protein ratios across individuals, including many rQTL that do not overlap with known individual protein-level QTL (i.e., pQTL) (Suhre, 2024). This important finding underscores that the ratio in the abundance of two proteins could be specifically regulated through shared genetic variance and non-genetic factors. Together, multiple lines of evidence now indicate that the ratiometric changes of protein pairs may present an important dimension to understanding fetal gene usage, one that has a tradition rooted in classic physiology studies but so far has not been broadly applied to unbiased discovery approaches.

Mass spectrometry is now routinely used to measure the levels of thousands of proteins in cardiac samples (Hasman et al., 2023; Karpov et al., 2024; Roberts et al., 2024; Wojtkiewicz et al., 2022). To our knowledge, existing proteomics studies have focused only on comparing the relative abundance of the same proteins across different conditions, and have not explicitly tested for the changes in ratios between proteins. One approach that can be taken to produce an unbiased catalog of protein ratios would be to test exhaustively for all pairwise protein pairs within a proteomics dataset. However, such an all-to-all ratio comparison would involve performing many statistical tests that could inflate family-wise error rates (e.g., a list of 2,000 proteins would require making ∼2 M comparisons). Instead, we choose to focus our attention on proteins that are most likely to share biologically significant regulations by prioritizing the comparison of isoforms, a term broadly defined to encompass members within a gene family that share significant sequence homology, often with overlapping functions and regulatory modalities. New proteins and isoforms can arise from: (1) two distinct genes that have diversified from gene or genome duplication events in the past, i.e., paralogs (Mantica and Irimia, 2025); or (2) variants within a single gene that arise from exon addition or omission, i.e., differential alternative splicing (Barbosa-Morais et al., 2012; Blencowe, 2017). Collectively, these two mechanisms have accounted for the creation of most new proteins, and provided the substrate for functional diversification within new cells or tissue types during metazoan evolution (Mantica et al., 2024; Mantica and Irimia, 2025). Both gene paralogs and splice isoforms are well established to play important roles in the gene regulation of fetal tissues. For example, fetal hemoglobin (HbF) is expressed in utero and shifts to adult hemoglobin (HbA) after birth in humans; whereas developmentally regulating splicing is known to affect cardiac genes in sarcomere, ion channels, and signaling (Jiang et al., 2024; Wang et al., 2016).

Here, we describe a new workflow to discover significant protein isoform changes from large-scale proteomics experiments, and apply it to characterize the fetal gene program of the heart. The workflow involves first acquiring absolute protein abundance estimates from quantitative mass spectrometry data, including those that utilize MS2-level tandem mass tag (TMT) multiplexing to compare multiple samples within a single experiment. This is followed by the extraction of log abundance ratios between isoform pairs while considering minor isoform ratio and % sequence identity, and statistical testing using linear models. The results recapitulate many known hallmark fetal genes in the heart, including myosin heavy chain (MYH6/MYH7), cardiac troponin (TNNI3/TNNI1), glucose transporter (GLUT1/GLUT4), enolase (ENO1/ENO3), and other glycolytic proteins. At the same time, we discover novel candidates for fetal gene isoforms, including a switch of usage among paralogs that function in mitochondrial and gene expression regulatory pathways. Extending the analysis to isoforms of the same gene, we find that alternative splicing derived protein isoforms (spliceoforms) provide a rich parallel source of the fetal gene program. Taken together, we present a catalog of the major detectable protein isoform changes in the fetal gene program of the mouse heart. We foresee this resource can complement individual gene and protein level investigations and provide insights in the studies of cardiac development, disease, and regeneration.

## Results

### Quantification of protein isoform ratios in fetal and perinatal hearts

Measuring the ratio between two proteins requires quantifying their abundance on the same scale, i.e., estimates of absolute abundance. However, a substantial portion of proteomics studies are performed using tandem mass tag (TMT) or similar MS2-based isobaric stable isotope multiplexing methods. Although MS2 multiplexing has the benefits of increasing scalability and reducing variance in quantitative comparisons, a drawback is that the reporter channel intensities reflect only the relative abundance of the same protein across different samples, and therefore do not directly lend themselves to comparison of two different proteins on an absolute scale (Zecha et al., 2019). To overcome this limitation, we implemented a proteomic ratio quantification workflow that works with MS2 multiplexed mass spectrometry data, by considering the combined MS1-MS2 intensity values of each identified peptide. The core intuition is that the precursor ion quantity of a peptide at the MS1 level can be distributed across multiple samples within a labeling block proportionally to each sample’s respective MS2 reporter channel intensity (Ahrné et al., 2015; Clark et al., 2019; H. Wang et al., 2023; J. Wang et al., 2023). Doing so allows us to take advantage of the multiplexing ability of MS2 isotope label based quantitation while being able to estimate the absolute protein quantity per sample needed to calculate protein ratios.

Accordingly, we generated TMT-labeled proteomics data from C57BL/6 fetal (E17) and postnatal (P1) hearts (n=5 each). Upon database search and quantification, we performed protein parsimony grouping and TMT isotopic purity correction. The MS1-level precursor chromatographic intensity and MS2-level TMT reporter intensity of each peptide were then extracted from the mass spectral data, which contained information of across-peptide intensities and across-sample intensities across experiments, respectively (**Figure 1A**). The resulting protein absolute quantification values from the composite MS1-MS2 data (**Table S1**) readily distinguish fetal and postnatal protein expression profiles in principal component analysis (**Figure 1B**); span over 5 orders of magnitude; and show strong overall agreement between fetal and postnatal hearts (**Figure 1C**); altogether demonstrating we are able to quantity both high and low abundance proteins consistently. <u>To validate the composite MS1-MS2 protein quantity</u> <u>values, we converted the values to protein copy numbers per cell, which allowed us to compare these</u> <u>values to aggregate absolute protein copy numbers in the mouse heart form multiple studies in the</u> <u>literature (PaxDB</u>) (Huang et al., 2023) (**Figure 1D**). Moreover, the overall reliability of the absolute protein estimates is reflected by a robust correlation with the estimated protein copy numbers in mouse fibroblasts from an independent study, despite differences in cell type and quantification methodology (**Figure 1E**). These results indicate the absolute protein abundance estimates are accurate and not negatively affected by MS2 multiplexing.

**Figure 1.**
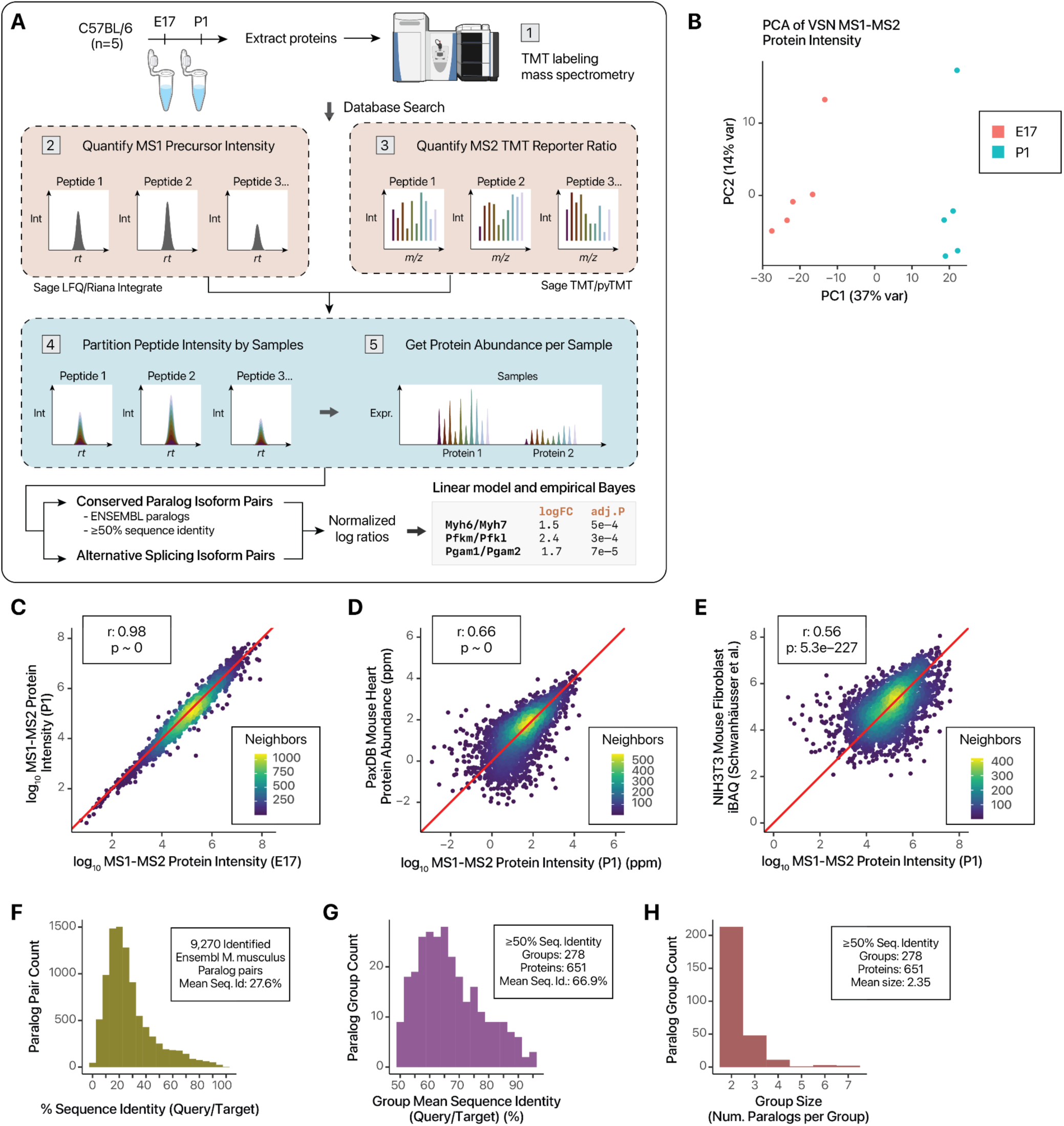
Experimental and analytical workflow of isoform usage tests. **A.** Schematic of workflow to perform protein absolute quantification from TMT data and apply limma to normalized log ratios. **B.** PCA from MS1-MS2 based protein absolute quantity clearly segregated fetal and postnatal heart samples. **C.** The derived protein absolute abundance values span over 5 orders of magnitude and are strongly correlated between E17 and P1 samples (Pearson’s r: 0.98, p ∼ 0 in log-log scale). Color denotes point density. **D.** The derived protein absolute abundance values are strongly correlated with known protein absolute abundance values in the mouse heart curated from multiple public data sets in PaxDB (Pearson’s r: 0.66, p ∼ 0 in log-log scale). **E.** The derived protein absolute abundance values are strongly correlated with mouse NIH3T3 fibroblast absolute data derived using iBAQ method in Schwanhäusser et al. 2011 (Pearson’s r: 0.56, p: 5.3e–227 in log-log scale). **F.** Histogram showing the distribution of sequence identity (query to target) of 9,270 Ensembl annotated M. musculus paralog pairs identified in the experiment (mean: 27.6%) **G.** Histogram showing the average sequence identity of paralog pairs in paralog groups selected for differential expression ratio analysis (≥50% sequence identity) (mean: 66.9%) **H.** Histogram showing the mean paralog group size among paralog pairs selected for differential expression ratio analysis (mean: 2.35)

After deriving the MS1-MS2 based absolute quantification values, we gathered the paralog groups within closely related gene families that were identified within the dataset by retrieving conserved gene paralog pairs using Ensembl genome annotation with additional filtering for at least 50% sequence identity, a conservative threshold chosen based on conventional cutoffs for average sequence homologies within a protein family, in contrast to more distant protein superfamilies (Finn et al., 2010). In total, 9,270 paralog pairs are identified (**Figure 1F**). After filtering for sequence homology total, the paralog groups consist of 545 quantifiable pairwise paralog relationships, with an average sequence identity of 66.9% (**Figure 1G**). Each paralog group contains an average of 2.3 proteins (**Figure 1H**). We then consider the pairwise generalized log ratios of the paralogs within each paralog group. Diagnostic plots confirm that the log ratios remain conforming to limma assumptions of normality and linearity (**Figure S1–S2**). A moderated t-test as implemented in limma is then performed on the differential abundance ratios of the 520 pairs of paralogs between fetal and postnatal mouse hearts (**Table S2**), out of which 142 pairs are significantly different at a conservative threshold of limma adjusted P < 0.01 and absolute log fold-change ≥ 0.5 (**Table S3**). To remove protein pairs where one isoform may be present only at very miniscule levels, we further introduce a second magnitude cutoff of minor isoform fraction ≥ 0.05, leading to a final set of 125 significant paralog pairs.

Before interpreting the ratio test results, we interrogate whether significant ratio differences cover proteins that are not differentially expressed at individual protein levels under identical significance thresholds. On an individual-protein level, we quantify the relative fetal/postnatal abundance of 3,889 canonical UniProt proteins, of which 1,025 are differentially expressed under identical significance thresholds (FDR-adjusted limma p value < 0.01 and absolute log fold-change ≥ 0.5) (**Table S4**). As expected, differentially expressed proteins are enriched in processes important for cardiac development including cell cycle, mRNA splicing, mitochondrial translation, and complex I biogenesis terms (gene set enrichment analysis GSEA FDR adjusted q-value < 0.01), indicating the data captures rich developmental stage specific differences in the proteomic landscape of fetal and postnatal hearts (**Figure S2**). Notably, out of 520 quantified paralog pairs, 115 pairs contain at least one protein that is not differentially expressed in single-protein comparison and 24 pairs had neither protein differentially expressed in single-protein comparison (**Table S5**). Hence, the paralog ratio comparisons are able to uncover additional information under uniform significance thresholds.

### Ratio comparisons capture known paralog usage in prenatal to postnatal shift

At the E17 stage in late gestation, the fetal mouse heart has undergone the majority morphogenetic development including chamber differentiation, valve formation, and cardiomyocyte expansion, and morphologically resembles postnatal hearts. At the same time, cardiac maturation continues to proceed including a gradual shift toward oxidative phosphorylation, while the retention of regenerative capacity distinguishes it from postnatal and adult hearts. The gene usage changes between E17 and P1 should therefore represent maturation and adaptation that are most relevant to the cardiac metabolism and functional aspects of the fetal heart program. We ask whether the paralog ratio data are able to recapitulate the differential usage of previously known features of the fetal genetic program. These hallmark fetal genes can be grouped into two major functional groups – those that affect contractile proteins and those that affect glucose metabolism (**Figure 2A**). Indeed, the proteomics data indicate that contractile proteins and glucose metabolism proteins showed the highest amplitudes in ratio shifts, consistent with them being the major components of the fetal heart program at E17 in contradistinction to P1 hearts (**Figure 2B**).

**Figure 2.**
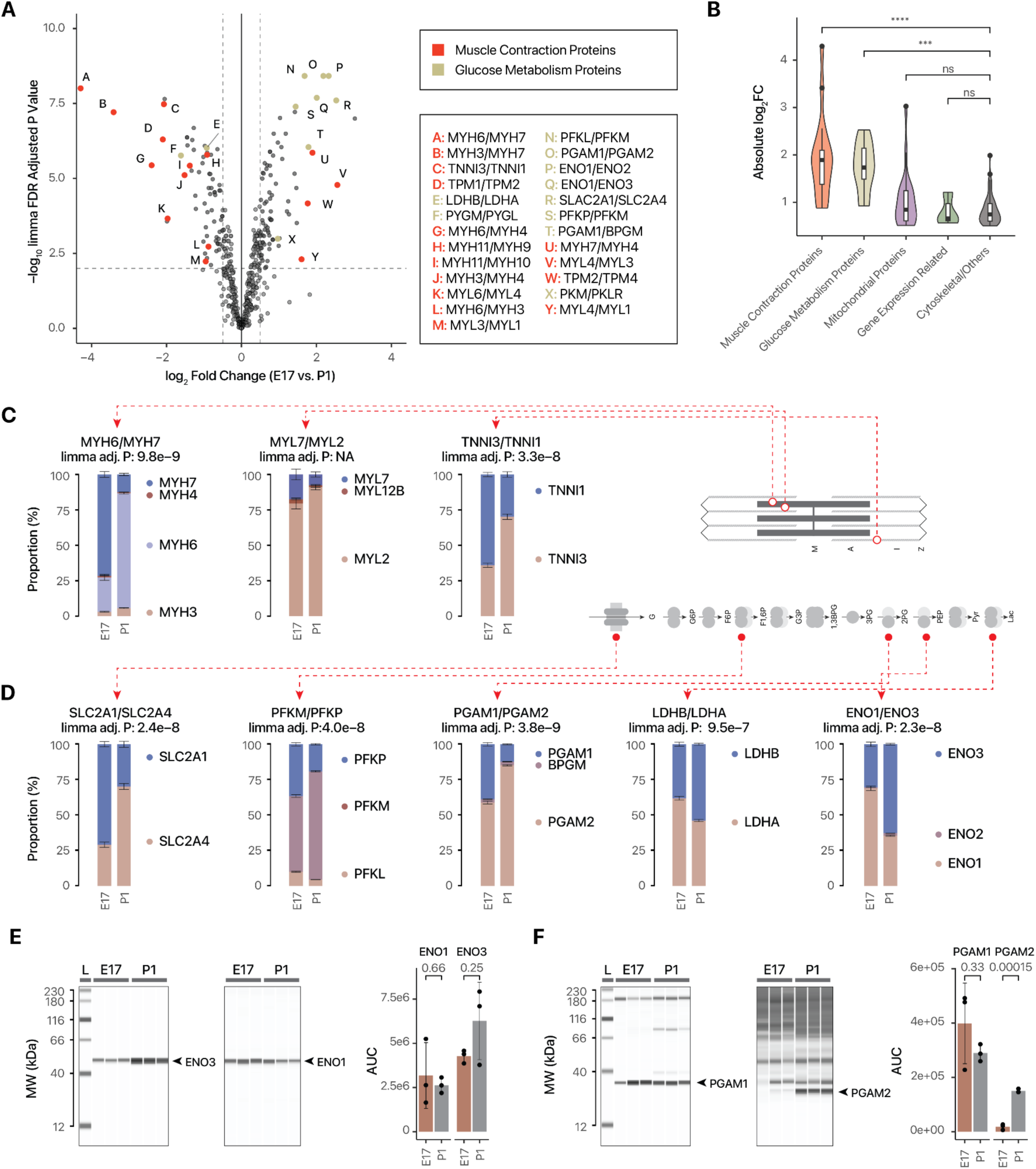
Protein isoform ratios capture hallmark fetal heart genes. **A.** Scatterplot of log2 fold-change (x-axis) vs. –log10 P value (y-axis) of compared paralog pairs (n=5 for fetal and postnatal hearts each) highlighting the significantly differentially expressed paralogs in contraction proteins (red) and glucose metabolism proteins (gold). Letters refer to paralog pairs in legends. **B.** Violin/boxplot of the magnitude of differential usage (absolute log_2_ ratio) of paralogs in different major functional categories between fetal and postnatal hearts. ****: two-tailed t-test P against “Cytoskeletal/Other” < 0.001; ***: P < 0.005; ns: p ≥ 0.05. **C.** Proportional bar charts showing the proportion of protein molecules in each paralog groups, reflecting the known shifts in isoform usage among sarcomere gene in the fetal gene program including (from left to right) MYH6/MYH7, MYL7/MYL2, and TNNI3/TNNI1. Error bars: s.e.m. of cumulative proportion. n=5 fetal and postnatal hearts. Paralog pairs are shown with limma adjusted P < 0.01, absolute logFC ≥ 0.5, and average minor isoform fraction ≥ 0.05, except for MYL7/MYL2 which was individually inspected as a known fetal gene program protein. **D.** Same as C, but for hallmark fetal genes in glucose metabolism, from left to right: SLC2A1/SLC2A4 (GLUT1/GLUT4), PFKM/PFKP, PGAM1/PGAM2, LDHB/LDHA, ENO1/ENO3. Paralog pairs are shown with limma adjusted P < 0.01 and average minor isoform fraction ≥ 0.05. **E.** Digital immunoblots after capillary electrophoresis corroborates the relative shift from ENO1 to ENO3 in postnatal hearts. P values: two-tailed t-test. Error bars: s.d. **F.** Same as E, but for the postnatal shift from PGAM1 to PGAM2.

#### (i) Changes in contractile protein isoform usage

The ɑ (MYH6) and β (MYH7) myosin heavy chain isoforms bind with a finite pool of actin and tropomyosin complexes in the sarcomere in different muscle fiber types and developmental stages. In the mouse, the β myosin heavy chain is a well-established hallmark of the fetal genetic program (Lyons et al., 1990; Reiser et al., 2001, p. 20). Consistent with this, the data clearly recapitulates a prominent ∼20-fold reciprocal shift from β to ɑ form from E17 to P1 hearts, where in the former MYH7/MYH6 ratio is >2 to 1, whereas in P1 hearts MYH6 is >8 fold higher (**Figure 2C**). The developmental stage specific expression of fetal troponin I is another well-established hallmark in fetal development (Bhavsar et al., 1991; Hunkeler et al., 1991). Likewise, the data reveal a >4-fold relative shift from slow skeletal muscle type troponin I (TNNI1) toward the adult cardiac type (TNNI3), where they are expressed ∼1:1 in E17 hearts but TNNI3 is >4 fold higher in P1 hearts (**Figure 2C**). In addition, a targeted query finds a suggestive change in MYL7 to MYL2, which reflects the known shift in MLC2a to MLC2v during cardiomyocyte maturation. Other changes in contractile proteins include the known difference in ventricular vs. fetal myosin light chains. The data recapitulates a >5 fold further enrichment of the ventricular MYL3 over fetal/essential MYL4 in postnatal hearts, where MYL3 is already expressed at ∼3-fold higher in the fetal heart than MYL4, but is dramatically enriched to ∼16-fold higher in postnatal heart. Another contractile protein usage difference concerns a 2.6-fold shift from the non-muscle myosin polypeptide MYH10 toward the smooth muscle myosin heavy polypeptide MYH11, where MYH10 is 2-fold higher than MYH11 in fetal heart, but lower in postnatal hearts (**Table S3**). Therefore, the proteome ratio test results readily capture well-established changes in sarcomeric isoform usage.

#### (ii) Changes in glucose metabolism protein isoform usage

Prominent changes are also seen among proteins that function in glucose metabolism. We consider here five known fetal heart genes. (1) First, fetal hearts express more basal glucose transporter GLUT1 (SLC2A1) than insulin-sensitive GLUT4 (SLC2A4) in a ∼70%/30% ratio, but roughly opposite proportion in postnatal hearts (30%/70% ratio) (**Figure 2D**). A transition from GLUT1 to GLUT4 in neonatal development is known in the heart and in brown fat of rats (Castelló et al., 1994), whereas prior work using transcript data whereas both GLUT1 and GLUT4 are more highly expressed in adult than fetal human hearts (Taegtmeyer et al., 2010). (2) Phosphofructokinase-1 (PFK-1; EC 2.7.1.11) is a critical glycolytic enzyme that catalyzes the committed step of fructose 6 phosphate to fructose 1-6, bis phosphate. PFK-1 exists as 3 isozymes encoded by 3 paralog genes: PFK-M (muscle); PFK-P (platelet); and PFK-L (liver); adult hearts are known to express exclusively PFK-M isoform. The ratio proteomics data recapitulates an expansion of the muscle isoform from fetal heart (53%) to postnatal heart (76%) coupled to the contraction of liver (10% to 4%) and platelet (37 to 19%) isoforms (**Figure 2D**), consistent with isozyme biochemical activity assays (Thrasher et al., 1981). (3) Another glycolytic enzyme, phosphoglycerate mutase (PGAM1/2; EC 5.4.2.11), the proportion of the skeletal muscle and myocardial enriched type-M isoform (PGAM2) increases from 59% to 85% of the paralog group between fetal to postnatal hearts, whereas the proportion of the type-B isoform PGAM1 decreases from 39% to 12% (**Figure 2D**). (4) In fetal hearts, alpha enolase (ENO1; EC:4.2.1.11) is the predominant isoform, amounting to 69% of abundance within the paralog group, but decreases to 35% in postnatal heart (**Figure 2D**). This is coupled with an increase from 30% to 63% of the striated muscle isoform of enolase (beta enolase, ENO3), a ∼4-fold switch in ENO1/ENO3 ratios. (5) In fetal hearts, the lactate dehydrogenase (EC:1.1.1.27) H and M subunits (LDHA)/(LDHB) exist at a 62%/38% ratio, as opposed to 46%/54% in postnatal hearts, an 1.9-fold switch (**Figure 2D**). This shift is consistent with the known enrichment of lactate dehydrogenase complexes containing the H subunit LDHB in the heart and its preference for conversion of lactate toward pyruvate in oxidative metabolism environment, contrary to the preference of the M subunit to favor lactate production from pyruvate in glycolytic tissues (Read et al., 2001). Other metabolic proteins include a decrease in proportion of non-muscle type cytosolic creatine kinase (CKB), but a reciprocal increase in the proportion of mitochondrial matrix creatine kinase (CKMT2) (**Table S3**). Therefore, the data captures well-established glycolytic gene usage differences.

#### (ii) Validation by orthogonal experiments and additional proteomics data

To corroborate the protein isoform ratios in fetal hearts, we adopt two orthogonal approaches. First, we note that although immunoblots can measure the relative fold-change of a protein between conditions, they cannot directly quantify the absolute proportion of two proteins without extensive calibration to map the measurements to absolute concentration units, due to the different affinity of antibodies to epitopes. Antibodies may also conflate closely related sequences. Nevertheless, these caveats notwithstanding, digital immunoblots following capillary electrophoresis (ProteinSimple Jess) using two specific antibodies against ENO1 and ENO3 are consistent with a postnatal toward ENO3 relative to ENO1 in E17 vs. P1 (**Figure 2E**). Likewise, immunoblots show a suggestive decrease in PGAM1 band intensity and a significant increase in PGAM2 band intensity, in agreement with the ratio proteomics data (**Figure 2F**).

As a second approach to immunoblots to further corroborate our findings, we apply the MS1-MS2 ratiometric calculation and statistical testing workflow to an existing TMT-labeled tandem mass spectrometry study containing proteomics time-series data during the development of embryonic (E10.5, E12.5, E14.5, E16.5, E18.5) and postnatal (P1, W1, W2, W4, W8) C57BL/6 mouse hearts (Gu et al., 2022). This analysis clearly reproduces hallmark sarcomeric and glycolytic fetal heart proteins, underscoring the consistency of ratio calculations across independent data sets (**Figure 3A**). Moreover, the isoform ratios of 299 common paralog pairs shared in both data sets are able to distinguish pre- and post-natal samples at different time points (**Figure 3B**), indicating strongly that paralog usage alone is sufficient to distinguish cardiac developmental stages. Taken together, we surmise that the proteomics ratio tests can capture numerous hallmark fetal genes at the protein level.

**Figure 3.**
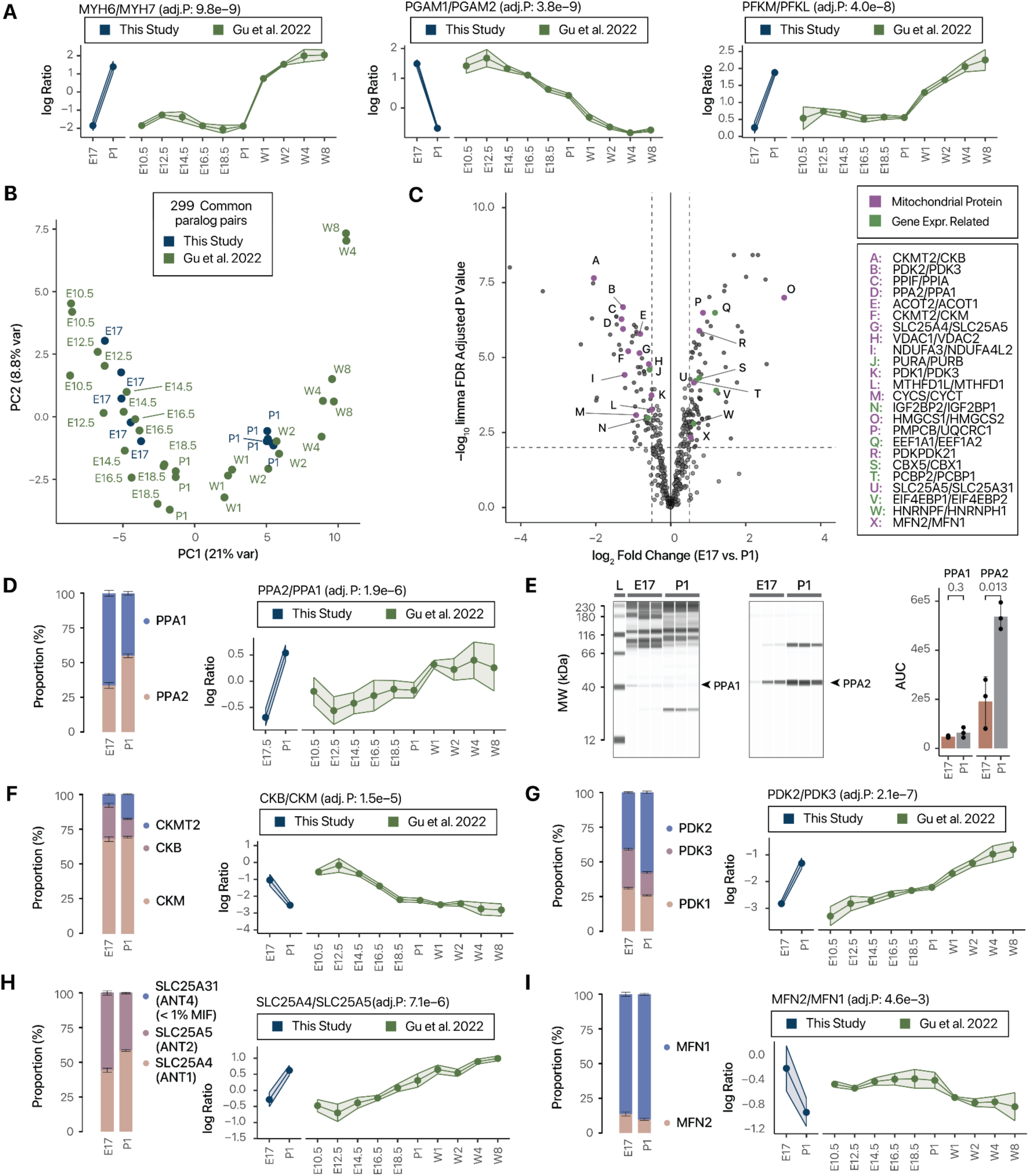
Detection and validation of paralog usage shifts in mitochondrial proteins. **A.** Application of ratiometric calculation to prior data recapitulated the rise in MYH6/MYH7, PGAM2/PGAM1, and PFKM/PFKL ratios in postnatal hearts across data sets. Adj. P: limma FDR adjusted P value in E17 vs. P1 (n=5) in this study. Error bar/ribbon width: s.d. **B.** PCA plot showing the abundance ratios of 299 commonly quantified paralog pairs distinguished developmental stages in the current study and re-analysis of an existing data set of pre- and postnatal C57BL/6 mouse hearts (Gu et al. 2022). **C.** Scatterplot of log2 fold-change (x-axis) vs. –log10 P value (y-axis) of compared paralog pairs (n=5 for E17 and P1 hearts each) highlighting the significantly differentially expressed paralogs in mitochondrial proteins (purple) and gene expression regulation related proteins (green). Letters refer to paralog pairs in legends. **D.** Proportional bar chart (left) showing an increase in PPA2/PPA1 ratio from E17 to P1 mouse heart (limma adjusted P: 1.9e–6), consistent with the postnatal increase of PPA2/PPA1 ratio in the re-analysis of Gu et al. 2022 (right). **E.** Digital immunoblots after capillary electrophoresis corroborates the relative shift from PPA1 to PPA2 in postnatal hearts. P values: two-tailed t-test. Error bars: s.d. **F.** Postnatal isoform usage shifts in CKB/CKMT across the two data sets. (Left) Bar chart error bars: s.e.m. of cumulative proportion. (Righ) ribbon chart error bar/ribbon width: s.d. **G.** Same as F, but for PDK2/PDK3. **H.** Same as F, but for SLC25A4/SLC25A5. **I.** Same as F, but for MFN1/MFN2.

### Ratio comparisons identify new candidate isoform shifts in fetal heart development

The results uncover a number of novel candidates of the fetal gene program. Most prominently, these proteins can be categorized as mitochondrial proteins and those functioning in gene expression regulation (**Figure 3C**).

#### (i) Mitochondrial protein isoform rewiring

We find changes in mitochondrial proteins including some not formally associated with the fetal gene program of the heart; for instance, a 2.4-fold reciprocal swing from inorganic diphosphatase PPA (EC:3.6.1.1) cytoplasmic isoform 1 (PPA1) to the mitochondrial isoform 2 (PPA2) (limma adjusted P 1.9e–6), from 66%/34% to 45%/55% (**Figure 3D**). This postnatal increase in PPA2 can be verified in the re-analysis of the Gu et al. 2022 data (**Figure 3D**) and experimentally using digital immunoblots (**Figure 3E**). Another mitochondrial related change in our data involves the relative shift from the cytosolic type-B creative kinase (EC 2.7.3.2) toward the type-M creatine kinase (CKM) polypeptide chain, in roughly 70%/25% to 70%/10% fashion (**Figure 3F**). The rest of the creatine kinase pool is accounted for by a postnatal increase in sarcomeric mitochondrial creatine kinase (smtCK) (CKMT2) which increases from less than 10% in fetal hearts to ∼20% of the pool in perinatal hearts, suggesting a reconfiguration of creatine kinase isozymes to increase phosphocreatine production for contractile demands. The decrease in CKB/CKM ratio is reaffirmed in the re-analysis of the Gu et al. 2022 time-series data. We note that these isoform switches cannot be fully explained by a simple increase in postnatal mitochondrial density, because wholly mitochondrial protein isoform groups also show differences in usage. For instance, we observe a switch from pyruvate dehydrogenase 1 and 3 (PDK1/PDK3) toward PDK2 as the major pyruvate dehydrogenase kinase isozyme (**Figure 3G**). All three isoforms are mitochondrial, suggesting the composition of mitochondrial proteins alters postnatally. In parallel, the cardiac skeletal muscle form of ADP/ATP carrier ANT1 (SLC25A4) replaces the fibroblast form (SLC25A5) ANT2 as the prevalent mitochondrial ADP/ATP carrier for ATP export (**Figure 3H**). This is accompanied by a further increase in MFN1/MFN2 ratios (**Figure 3I**) and a further increase of the ratio of VDAC1/VDAC2 ratio in the mitochondrial voltage-dependent anion channel (**Table S3**).

Other proteins relevant to mitochondrial biology include a shift of sideroflexin, a mitochondrial membrane transporter involved in heme metabolism and serine transport (Acoba et al., 2021; Kory et al., 2018), from the ubiquitous SFXN1 isoform toward the neuron-enriched SFXN3 isoform, which share close homology (77%) (**Table S3**). In parallel, hydroxymethylglutaryl-CoA (HMG-CoA) synthase (EC 2.3.3.10) HMGCS1 cytosolic isoform decreases, coupled to an increase in the mitochondrial form HMGCS2, leading to a strong switch toward HMGCS2 usage (logFC 3.03, limma adjusted P 1.1e–7), such that the proportion of HMGCS1 drops from 14% of the total HMGCS pool at E17 to 2% at P1. An HMGCS1 decrease during prenatal cardiac development is noted in previous work as part of the rewiring of the mevalonate pathway for cholesterol biosynthesis (Edwards et al., 2023). The data here in turns reveals a parallel increase in the mitochondrial form in early postnatal hearts.

During fetal cardiac development and toward birth, there is a gradual shift from a more glycolytic preference toward aerobic metabolism in the heart, which involves mitochondrial biogenesis as well as other processes (Zhao et al., 2019). Taken together, the observations here suggest that while the amplitude of ratio shifts in these proteins are smaller than those in sarcomeric and glycolytic proteins, metabolic remodeling of the postnatal heart may also involve broad parallel changes in isoform usage in mitochondrial proteins in addition to changes in glycolytic proteins.

#### (ii) Post-transcriptional and translational gene regulation programs

Changes related to gene regulation and structural proteins include a switch in the chromobox protein homologs CBX5/CBX1; the translation elongation factor alpha (EEF1A) from the “ubiquitous” EEF1A1 form toward the heart/muscle form EFF1A2, where the former is 6.5-fold higher in fetal hearts but about 3-fold higher in postnatal P1 hears and decreases further in later time points; and the poly(rC)-binding proteins PCBP1/PCBP2 (**Figure 4A**). To explore the potential functional implications of isoform usage shifts, we investigated the effect of PCBP1/PCBP2 expression in human AC16 cardiac cells. PCBP1/PCBP2 are RNA binding genes that contain KH RNA-binding domains, and are multifunctional proteins that have been implicated in transcriptional (Karam et al., 2024), post-transcriptional (Ishii et al., 2020) and translational regulations (Fujimura et al., 2008). How PCBP1 and PCBP2 are functionally differentiated remains a subject of interest (Ishii et al., 2020), and thus far there have been few reports on their function in cardiac cells. The two paralogs share close homology with ∼85% identical sequences; moreover, they share partially overlapping RNA binding targets including PCBP2 itself from eCLIP data, suggesting they may form mutually regulatory relationships. Human and mouse PCBP1 sequences share 100.0% identity, whereas human and mouse PCBP2 are identical except in 5 positions; therefore we expressed the human/mouse PCBP1 and human PCBP2 coding sequences in human AC16 cardiac cells. Immunofluorescence imaging confirms broad cytonuclear localization of both paralogs under exogenous expression (**Figure 4B**). RNA sequencing experiments of cells overexpressing PCBP1 or PCBP2 reveal the differential expression of 1362 and 672 transcripts, respectively (shrinkage s-value < 0.05; abs(logFC) > 0.25), including known eCLIP targets as well as those other genes, suggesting the differential expression may arise from both direct RBP target regulations and secondary effects (**Figure 4C**). A comparison of differential gene expression between PCBP1 and PCBP2 shows a high concordance (Pearson’s r 0.59 in log-log scale; p ∼ 0) which is consistent with the two proteins having overlapping roles on functional phenotypes (**Figure 4D**). Nevertheless, a small number of off-diagonal data points offer evidence of protein-specific effects and indicate the two paralogs show functional specialization by influencing the expression of non-overlapping genes (**Figure 4D**). Notably, some differential expression events including F4, HCFC2, GADD45A, RSRP1, LIF, and ATF3 appear to be partially neutralized when both isoforms are overexpressed together in the same cells (**Figure 4E**), which hint at the notion that the differential status (i.e., ratio) rather than the individual abundance of the PCBP1/PCBP2 proteins could be a determinant in functional outcome.

**Figure 4.**
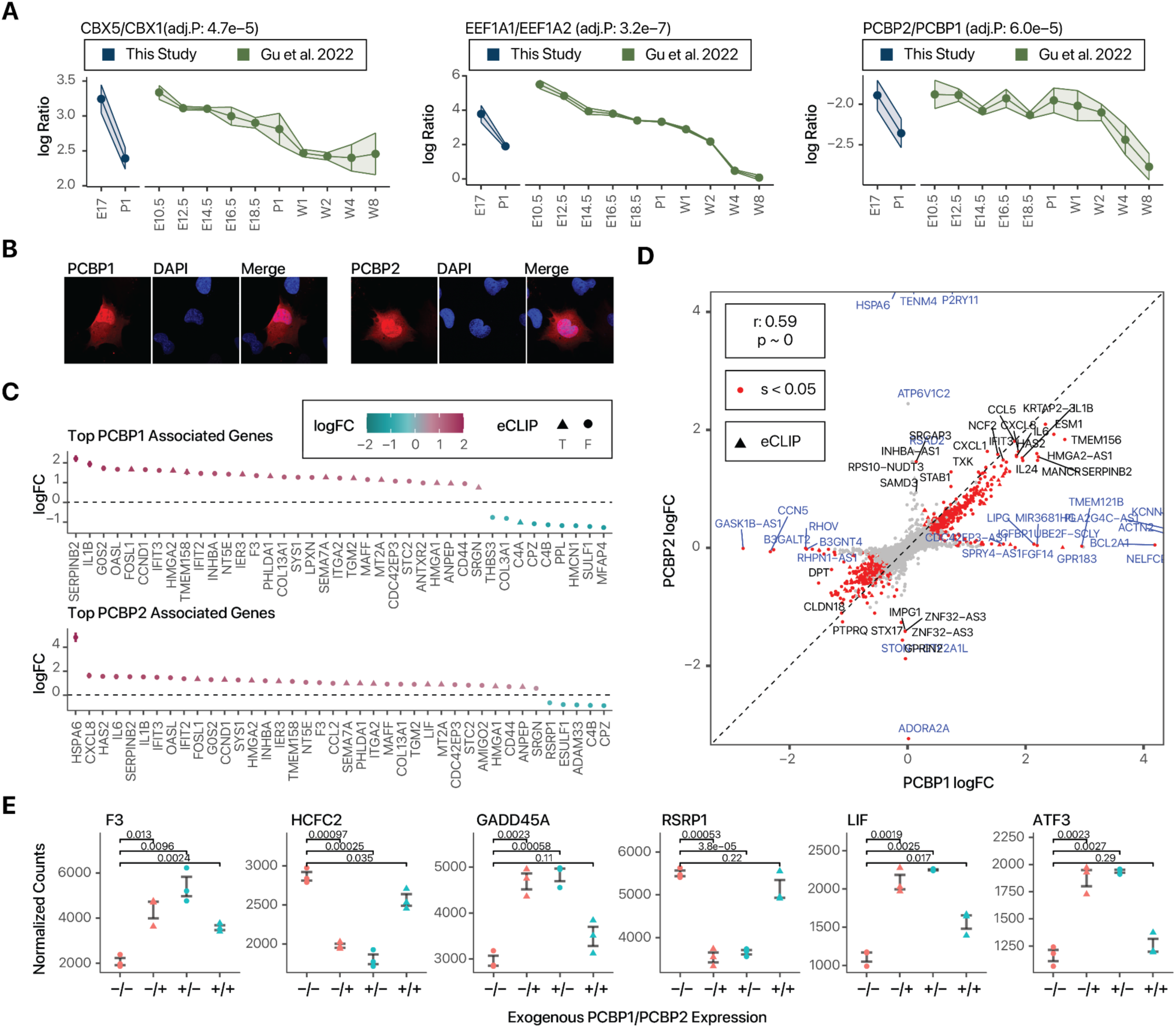
Examples of isoform usage shifts in gene regulation related proteins. **A.** Postnatal shifts in the ratio of gene expression regulation related proteins in data generated in this study and in the re-analysis of Gu et al. 2022. Adj. P: limma FDR adjusted P value in E17 vs. P1 (n=5) in this study. Error bar/ribbon width: s.d. **B.** Immunofluorescence images of human AC16 cardiac cells overexpressing HaloTag linked PCBP1 and PCBP2, showing nucleocytoplasmic localization of both proteins. **C.** RNA sequencing data showing the top differentially expressed genes associated with the expression of exogenous PCBP1 (top) and PCBP2 (bottom) in human AC16 cells. Colors denote log2 fold-change vs. control cells. Data point shape of triangle denotes that the gene is a known binding target of PCBP1 or PCBP2 in each respective plot in ENCODE eCLIP data. **D.** Scatterplot showing the overall similarities in the RNA abundance profiles of AC16 cells in PCBP1 (x-axis) and PCBP2 (y-axis) over-expression over control cells (Pearson’s correlation *r*: 0.59, p ∼ 0 in log-log scale). Red data points denote significantly differentially expressed genes (DESeq2 shrinkage s value < 0.05). Blue labels denote genes that show different regulations (off-diagonal genes) between PCBP1 and PCBP2. **E.** Dot plots showing the effects of PCBP1 and/or PCBP2 over-expression on the abundance (normalized counts) of select genes that show an interaction effect. –/–: no overexpression; –/+: PCBP1 overexpression only; +/–: PCBP2 overexpression only; +/+: concurrent PCBP1 and PCBP2 overexpression in the same cells.

### (iii) Other proteins

Protein switches that do not fall into the above categories include a switch from DYNLL1 toward DYNLL2 as the major isoform in postnatal hearts (50%/50% in fetal hearts, vs. 32%/68% in postnatal hearts); a subtle switch of vimentin (VIM)/desmin (DES) ratios (76%/24% to 64%/35%); a slight increase in lamin B2 (LMNB2) vs B1 (LMNB1) (**Table S3**); and a ∼2-fold further enrichment of fatty acid binding protein FABP3 over FABP4 (from 60/40 ratios to 75/25 ratios) in postnatal hearts (**Table S3**), which was also previously observed in single-cell transcripts in developing mouse hearts from E9.5 to P21 (DeLaughter et al., 2016). Taken together, the results demonstrate that the data and ratio calculation can be applied to analyze the temporal expression of protein isoforms during cardiac development to discover both known and novel isoform shifts. The reliability of the protein ratio analysis is underscored by the rediscovery of classic hallmark fetal gene program proteins, validation by external mass spectrometry data sets, and corroboration by orthogonal immunobiological experiments.

### Fetal to postnatal isoform shifts are enriched in evolutionarily recent paralogs

We next examined the relationship between the differential usage of isoform pairs and their evolutionary conservation. Consistent with known ancestry of gene families, the majority of quantified paralogs belong to families that are ancient (inferred to originate in or before the common ancestors of all bilaterians) (**Figure 5A; Table S6**). Also as would be expected, more recent paralogs share greater sequence homology as defined by the mean (query-target and target-query) sequence identity following ClustalO sequence alignment (**Figure 5B**). We notice a modest trend for higher average sequence identity of a pair of paralogs with whether it is differentially expressed in fetal vs. postnatal hearts, although this trend is not statistically significant (two-tailed t test P: 0.42) (**Figure 5C**). On the other hand, we observe a statistically significant relationship showing that more ancestral paralog pairs, such as those that are conserved across opisthokonta lineages, are less likely to be differentially expressed in fetal vs. postnatal hearts (Fisher exact test P: 0.021) (**Figure 5D**). This statistically significant relationship can also be seen when comparing Ensembl paralog types, which distinguish ancient paralogs as those inferred indirectly from supertrees, again showing ancient paralogs are less likely to be differentially expressed (Fisher’s exact test P: 6.5e–3) (**Figure 5E**). This suggests that the fetal heart program may compose preferentially of evolutionarily more recent paralogs.

**Figure 5.**
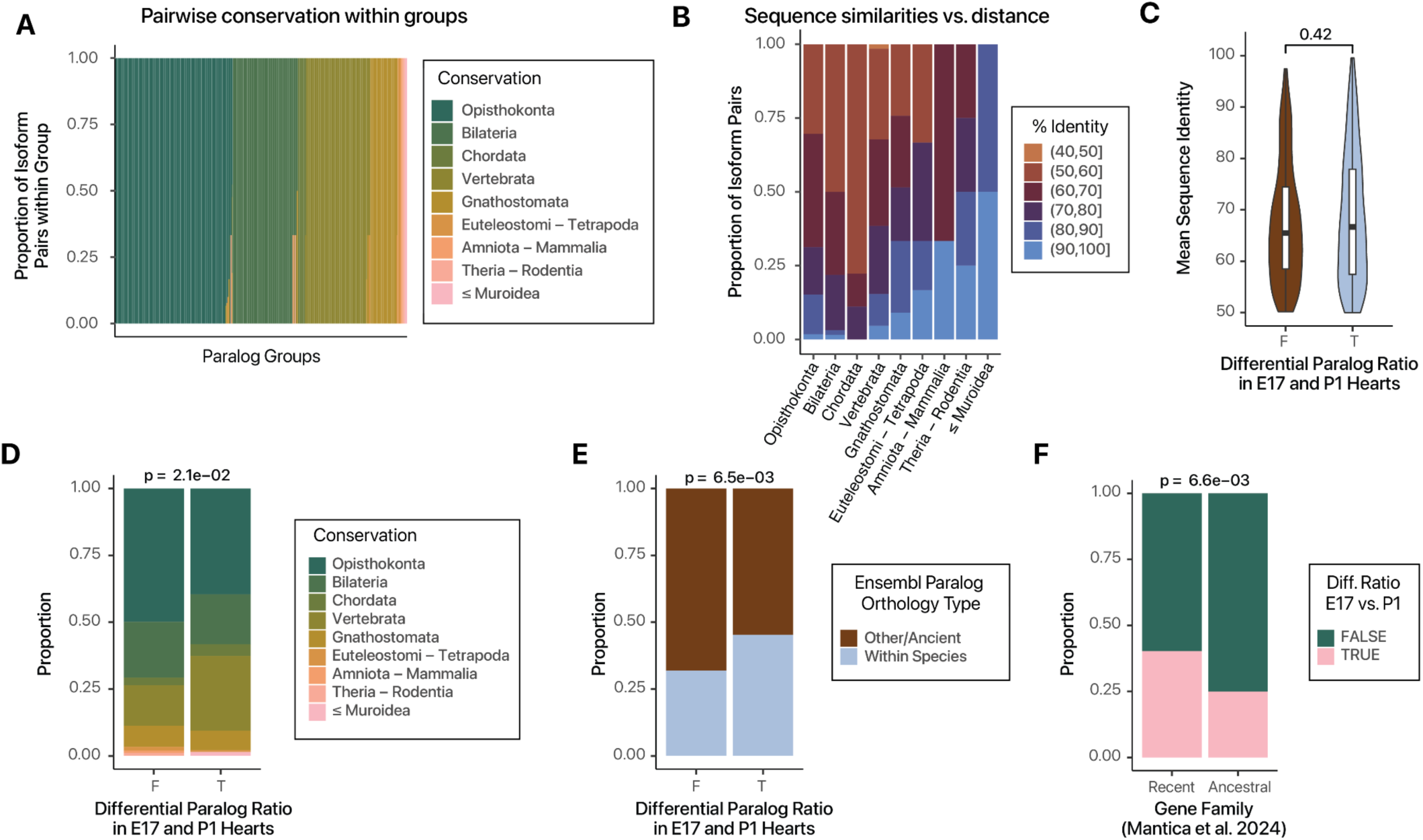
Evolutionary conservation of fetal regulated paralogs. **F.** Sorted proportional bar charts showing the evolutionary conservation of all paralog pairs within each of the paralog groups from the most deeply conserved (across Opisthokonta) to the least (Muroidea to Mus). **G.** Proportional bar charts showing the binned average sequence homology among all paralog pairs within each conservation distance category. Open bracket “(“: non-inclusive interval; closed bracket “]”: inclusive interval. **H.** Violin/box plots of paralog differential expression status in fetal hearts vs. mean sequence identity (query-target). P value: two-tailed t-test. **I.** Proportional bar charts showing the ancestry distribution of paralog pairs that have consistent vs. differential expression ratios in E17 vs. P1 hearts. P values: Fisher’s exact test for Opisthokonta membership. **J.** Proportional bar charts showing Ensembl paralog type vs. differential expression ratios in E17 vs. P1 hearts. P values: Fisher’s exact test. **K.** Proportional bar charts showing ancient vs. recent gene family categorization of paralog pairs (Mantica et al. 2024) against differential expression ratios in E17 vs. P1 hearts. P values: Fisher’s exact test.

To corroborate this observation, we compared fetal expression specialization with evolutionary conservation in bilateria, using the results of a recent study that reconstructed the phylogeny of gene expression profiles from 20 species to infer gene ancestry (Mantica et al., 2024). By considering gene families that originated from or predated the common ancestor of bilaterians to be ancient and those within more recent families to be recent (Mantica and Irimia, 2025), the results likewise indicate that evolutionarily recent genes are significantly more likely to be regulated in fetal expression (Fisher’s exact test P: 6.6e–3) (**Figure 5F**). Taken together, these analyses show that evolutionarily more recent paralogs are enriched in expression specialization in fetal hearts, which is consistent with the notion of continued functional specialization and neofunctionalization of paralogs during metazoan evolution.

### Spliceoforms compose a second source of fetal isoform ratio shifts in the heart

Alternative splicing presents a second major mechanism to generate gene isoforms. At present, the identification of splicing-derived isoform proteins (i.e., spliceoforms) remains highly challenging, due to both experimental and data analytical reasons (Blencowe, 2017; Lau et al., 2019; Wang et al., 2018). For instance, alternative isoforms are often found in much lower abundance than their canonical counterpart. Compounded to this, splice junctions between exons are known to preferentially code for lysines, which become cleaved by trypsin in typical proteomics workflows, leading to the loss of exon-exon connection information in mass spectrometry experiments. Moreover, many isoforms are tissue specific and may not be catalogued in commonly employed protein sequence databases, which prevents their identification by database search algorithms.

To examine whether spliceoform ratios also differ between fetal and postnatal hearts, we first searched the data against mouse UniProt/Swiss-Prot canonical and isoform sequences. Applying the same protein ratio calculation and significance threshold as the paralog pair comparisons (FDR adjusted P < 0.01, ≥ 5% minor isoform fraction, absolute log_2_ fold-change ≥ 0.5), we find 11 pairs of isoforms whose ratios differ significantly between E17 and P1 hearts. To help confirm the isoform identifications, we further use a second search engine and protein quantification approach to validate the identification, which is an RNA-seq guided workflow that comprises JCAST, MSFragger, Riana, and pyTMT (see Methods). Among shared proteins, both workflows return highly similar protein absolute values over 5 orders of magnitude (correlation coefficient: 0.85 – 0.86) (**Figure S4**). This second workflow is able to confirm 10 of the 11 significant isoform shifts found in our main search strategy (**Figure 6A**) and further uncovers 9 additional significant isoform pairs at adjusted P < 0.01, minor isoform fraction > 0.05, bringing the total number of protein splice isoform signatures to 20 (**Table S7**).

**Figure 6.**
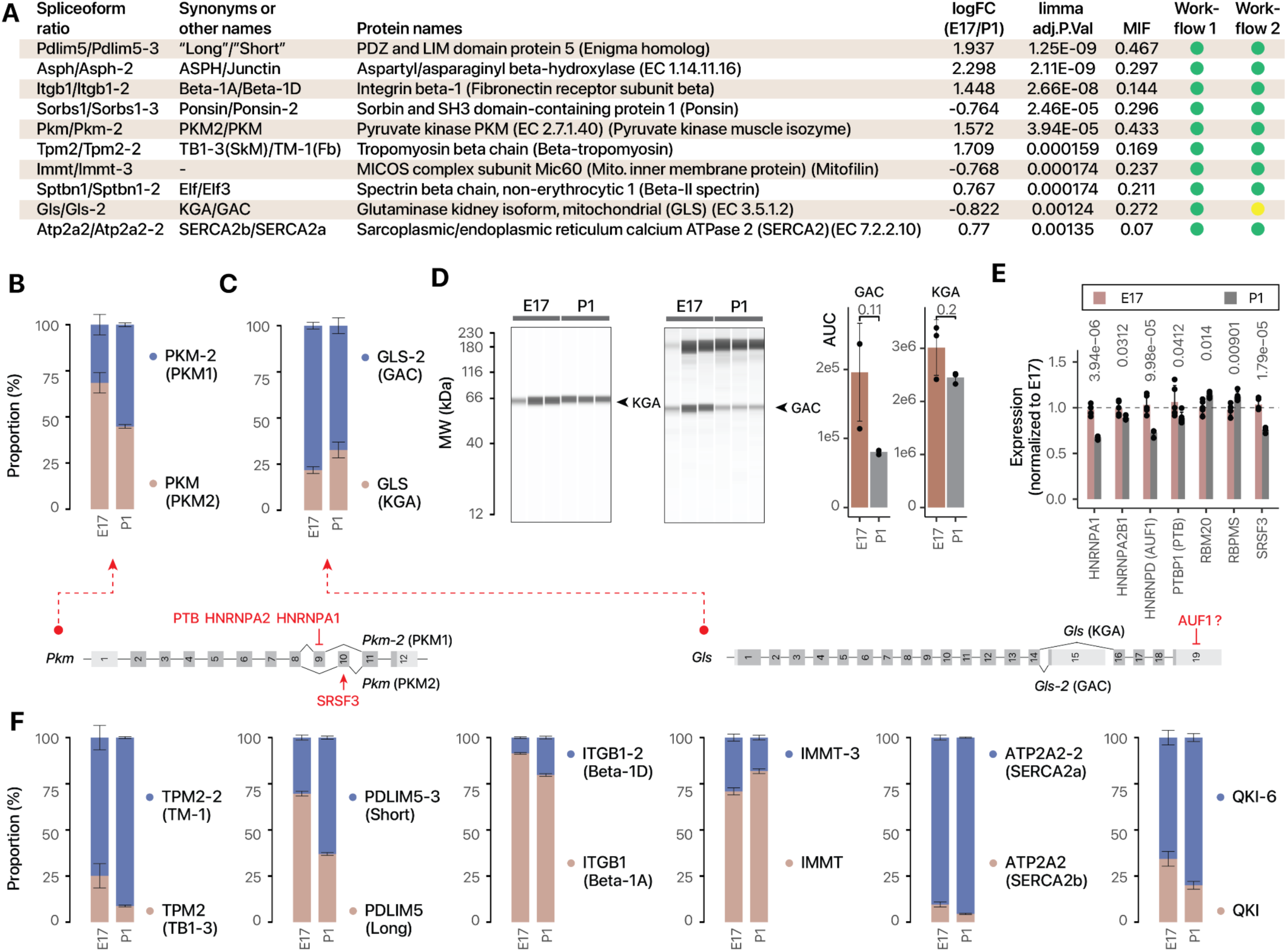
Alternative splicing derived protein isoforms as a source of fetal genes. **A.** The names and synonyms of 10 spliceoform pairs that are significantly different between E17 vs. P1 hearts in both database search/quantification workflows employed. The values of limma FDR adjusted P value (adj.P.Val) and log2 fold-changes (logFC) refer to results of Workflow 1 (Sage against UniProt/Swiss-Prot isoforms). Workflow 2 refers to MSFragger/Riana/pyTMT against a JCAST-generated tissue-specific protein sequence database. Green: FDR ≤ 0.01, absolute logFC ≥ 0.5; minor isoform fraction (MIF) ≥ 0.05. Yellow: FDR ≤ 0.01, minor isoform fraction (MIF) ≥ 0.05. **B.** Proportional bar charts showing the estimated proportions of PKM splice isoforms (M1 and M2), reflecting the known shifts from M2 to M1 in postnatal hearts. Error bars: s.e.m. of cumulative proportion. n=5 fetal and postnatal hearts. **C.** Same as B, but for the GS isoforms KGA and GAC. **D.** Digital immunoblots after capillary electrophoresis corroborates the relative shift from GAC to KGA in postnatal hearts. P values: two-tailed t-test. Error bars: s.d. **E.** Bar charts showing the normalized (to E17) expression of splice factors and RNA binding proteins implicated in isoform shifts. Numbers denote limma FDR adjusted P values in E17 vs. P1 hearts. **F.** Proportional bar charts for additional splice isoform pairs TPM2 (TM-1 vs. TB1-3), PDLIM5 (long vs. short), ITGB1 (beta-1D vs. beta-1A), IMMT (canonical vs. -3), ATP2A2 (SERCA2a vs. SERCA2b) and QKI (canonical vs. -6). Error bars: s.e.m. of cumulative proportion. n=5 fetal and postnatal hearts. Selected spliceoform pairs are shown with limma adjusted P < 0.01; average minor isoform fraction ≥ 0.05.

Pyruvate kinase (EC 2.7.1.40) has two main splice isoforms in the heart: the adult M1 form and the fetal M2 form (n.b. The M2 isoform has been designated as the canonical entry on UniProt and Ensembl, hence the M1 isoform’s assigned database name in this study is PKM-2) (**Figure 6B**). The M2 isoform differs from M1 by a pair of mutually exclusive exons at exon 9 (M1) and exon 10 (M2) and is known to be reactivated in failing hearts (Rees et al., 2015). The protein ratio results show the expected switch of usage from PKM2 to PKM1 in P1 hearts. Glutaminase 1 (GLS-1; EC 3.5.1.2) is a main rate-limiting enzyme in glutaminolysis, and switches from the high-activity GAC to the KGA isoform (**Figure 6C**). This switch can be partially corroborated by immunoblots, despite some limitations of the approach as stated above (**Figure 6D**). Upregulation of glutaminolysis has been implicated in cardiac hypertrophy and failure (Yoshikawa et al., 2022), suggesting GLS-1 may form a part of the constellation of fetal metabolism genes with differential usage during cardiac development and remodeling.

These instances of differential isoform usage are underpinned by concurrent RNA binding protein (RBP) regulations in postnatal hearts. The splicing of PKM2 is known to be controlled mechanistically by three splice factors, HNRNPA1, HNRNPA2, and PTB; which function together to suppress the inclusion of the M1-specific exon 9 (Israelsen and Vander Heiden, 2015). Another splice factor, SRSF3, promotes the exclusion of the M2-specific exon 10. Consistent with this, we see that these splice factors are preferentially expressed in E17 hearts and depleted in P1, which would elevate PKM2 expression in fetal hearts (**Figure 6E**). The regulation of KGA and GAC isoform usage is more complex and likely involves multiple RBPs, lncRNAs, and protein degradation factors (Adamoski et al., 2024; Masamha et al., 2016). In particular, HNRNPD (AUF1) is known to bind AU-rich elements at AUUUA pentamers (White et al., 2017), which are present in the respective 3’ UTR of both GAC and KGA transcripts. Prior work studying the gene sequences in the rat suggests that AUF1 may destabilize KGA (Masamha et al., 2016). In our hands, AUF1 is repressed in P1 hearts (limma adjusted P: 1e–4) (**Figure 6E**), suggesting it may play a relevant role in the postnatal isoform shift.

Mammals have four tropomyosin genes: TPM1 (alpha), TPM2 (beta), TPM3, and TPM4. Moreover, each of four genes also has splicing-derived isoforms. Cardiac muscles predominantly express alpha-tropomyosin TPM1; whereas TPM2 beta-tropomyosin is a minor isoform in the adult heart, and is primarily associated with the embryonic heart. TPM2 has at least two splice variants: a skeletal muscle form (TB1-3) that includes exon 6B (also referred to as exon 7) and exon 9A; and a smooth muscle isoform (TM-1) that includes exon 6 (also referred to as 6A) and 9D. In the postnatal heart, beta-tropomyosin expression is known to decrease, which is seen expectedly in the data. Moreover, we observe that this decrease primarily affects the skeletal muscle isoform TB1-3 (**Figure 6F**). This observation is corroborated by previous RNA-seq data, which shows that in wild-type hearts, the smooth muscle isoform TM-1 is predominantly expressed over the skeletal muscle isoform (Montañés-Agudo et al., 2023). The mutually exclusive exon pairs 6 and 7 (6A and 6B) are known from targeted studies to be regulated by PTB, which mainly serves to suppress exon 7 usage (Mulligan et al., 1992; Ye et al., 2013). This may seem incongruent given the postnatal decrease of PTBP1 in the heart. A potential explanation may be found from the meta-analysis of splice factor knockout experiments, which revealed that TPM2 splicing requires the combined action of multiple splice factors, including RBM20, which suppresses exon 7/6B (Montañés-Agudo et al., 2023). Indeed, we observe a postnatal increase in RBM20 (limma adjusted P 0.014), that may contribute to the decrease in the skeletal muscle TPM2-specific exons (**Figure 6E**).

Another spliceoform shift involves PDLIM5 (**Figure 6F**), which switches from the canonical form toward the PDLIM5-3 sequence. PDLIM5 has several splice variants that have been referred to as the “long” and “short” isoform groups in the literature. PDLIM5-3 is one of the two shorter sequences of mouse PDLIM5 that are documented on Swiss-Prot, and misses residues 338–591 in the canonical PDLIM5 sequence. It therefore corresponds to a “short” PDLIM5 isoform and may correspond to the variant that is known to participate in cardiomyocyte binucleation during cardiac development (Gan et al., 2022). Paradoxically, whereas prior work has found that RBPMS deletion leads to the accumulation of the short isoform, we find here that the postnatal increase in the short PDLIM5-3 spliceoform is associated with a slight but significant increase in RBPMS levels in postnatal hearts (**Figure 6E**), suggesting other factors may also influence PDLIM5 splicing. Yet other spiceoform changes include the alternative splicing of integrin beta 1, causing a relative expansion of the striated muscle 1D isoform (ITGB1-2) (Belkin et al., 1996, p. 1) over the canonical 1A (ITGB1) isoform (**Figure 6F**). Mitofilin (IMMT), which forms part of the MICOS complex in the mitochondrial inner membrane, shows a depletion of the uncharacterized -3 isoform in postnatal hearts (**Figure 6F**). The splicing of IMMT may also be influenced by RBM20 activity (Kornienko et al., 2023). In parallel, we see a shift from SERCA2b (ubiquitous isoform) toward SERCA2a (muscle specific) (**Figure 6F**). While the SERCA2a spliceoform is known to be specific to slow twitch skeletal and cardiac muscle, the results here provide further evidence that its differential splicing may form part of the fetal gene program.

Finally, using the JCAST/MSFragger workflow (Workflow 2), we further identify an isoform shift in QKI (Quaking, KH domain containing RNA binding) (**Figure 3G**); QKI is a cardiac RBP that regulates alternative splicing and is important in processes ranging from cardiomyocyte differentiation to ischemic injury. A re-analysis of large-scale data shows that QKI regulates the splicing of many important cardiac genes, including calmodulin-dependent protein kinase CAMK2D, beta-tropomyosin TPM2, and titin (TTN) (Montañés-Agudo et al., 2023). QKI itself has three major splice isoforms: the designated canonical QKI-5, and the alternative isoforms QKI-6 and QKI-7 with different C-terminus residues. We find an increase in QKI-6 over QKI-5 ratio in the postnatal heart, which mirrors the decline of QKI-5 that is seen over embryogenesis in the brain (Larocque et al., 2009). Whereas QKI-5 has a nuclear localization signal and is thought to regulate pre-mRNA splicing, QKI-6 is largely cytoplasmic (Larocque et al., 2009) and may function in RNA stability or post-transcriptional regulations. In the heart, deletion of *QKI* locus leads to abrogated cardiomyocyte contractility in hESC that is rescued by canonical QKI or (QKI-5) but not QKI-6, further supporting their functional diversification in cardiac cells (Chen et al., 2021). A change in QKI-5 and QKI-6 ratios from the same gene may therefore signify in a change RNA-binding function that together with other known RBP changes in cardiac development (D’Antonio et al., 2022; Gan et al., 2022) can initiate a broad reordering of splicing networks. Taken together, although the analysis underscores the continued challenges in identifying spliceoform proteins, we can now readily quantify a number of known and novel alternative splicing isoform switches. The results here further show that spliceoform switches provide a rich source of fetal gene candidates that call for further investigations.

### Fetal isoform ratios are partially reversed in pathological cardiac remodeling

We next asked whether the isoform ratio shifts are reversed in pathological cardiac remodeling. To do so, we examined adult and aged mice, administered with saline or 20 mg/kg isoproterenol for up to 14 days to induce pathological cardiac hypertrophy (**Figure 7A–D**). On an individual protein and pathway levels, the isoproterenol treatment led to drastic increases in striated muscle contraction proteins and the depletion of mitochondrial metabolism and respiratory chain proteins, showing the experiments accurately capture the proteomic landscape of pathological hypertrophy (**Figure S5**; **Table S8**). We then focus on which isoform pairs are significantly different (limma FDR adjusted P value ≤ 0.1) in hypertrophic hearts that show the same direction of changes in fetal over postnatal comparisons, regardless of their log fold-change (**Table S9**). Under the applied threshold, we find several glycolytic proteins that show significant changes in isoform usage in hypertrophy and partially revert the direction of the fetal gene profiles (**Figure 7E**). Prominently, this includes a return to a lower LDHB/LDHA ratio. Overexpression of LDHA in neonatal rat ventricular myocytes has been shown to drive hypertrophic cardiac growth in rodent models (Dai et al., 2020), suggesting the possibility that this particular isoform shift may contribute maladaptively to hypertrophy. In addition, we observe a return toward a lower PFKP/PFKM ratio, the latter of which is known to be up-regulated in failing human hearts and may be maladaptive (Vigil-Garcia et al., 2021). Another glycolytic protein PGAM1/2, likewise shows a return toward a higher PGAM1/PGAM2 ratio. Additionally, We observe a return toward a lower inorganic phosphatase isoform PPA2/PPA1 ratio, which are important metabolic proteins in the mitochondria but have not been explored in the context of cardiac development or remodeling. There is a return toward a higher EEF1A1/EEF1A2 ratio, two homologous members of the eEF1A component of the eEF1 complex with unclear distinction of function. In each case however, only a partial reversal was observed and moreover, the magnitude of changes in hypertrophy appeared smaller than reported transcript changes (**Figure 7E**). To corroborate this observation, we applied the protein ratio calculation to a set of label-free mass spectrometry data generated by Arrell et al. (Arrell et al., 2020) who performed LAD coronary artery ligation to induce myocardial infarction (MI) in mice then measured cardiac remodeling after 4 weeks. We likewise observe a partial reversion to the fetal program in PGAM1/2 and PDK1/2 that is less in amplitude than the fetal/perinatal change (**Figure 7F**). This result suggests the partial nature of fetal gene reactivation at the protein level cannot be fully explained by the isoproterenol model or the ratio compression effect of TMT labeling.

**Figure 7.**
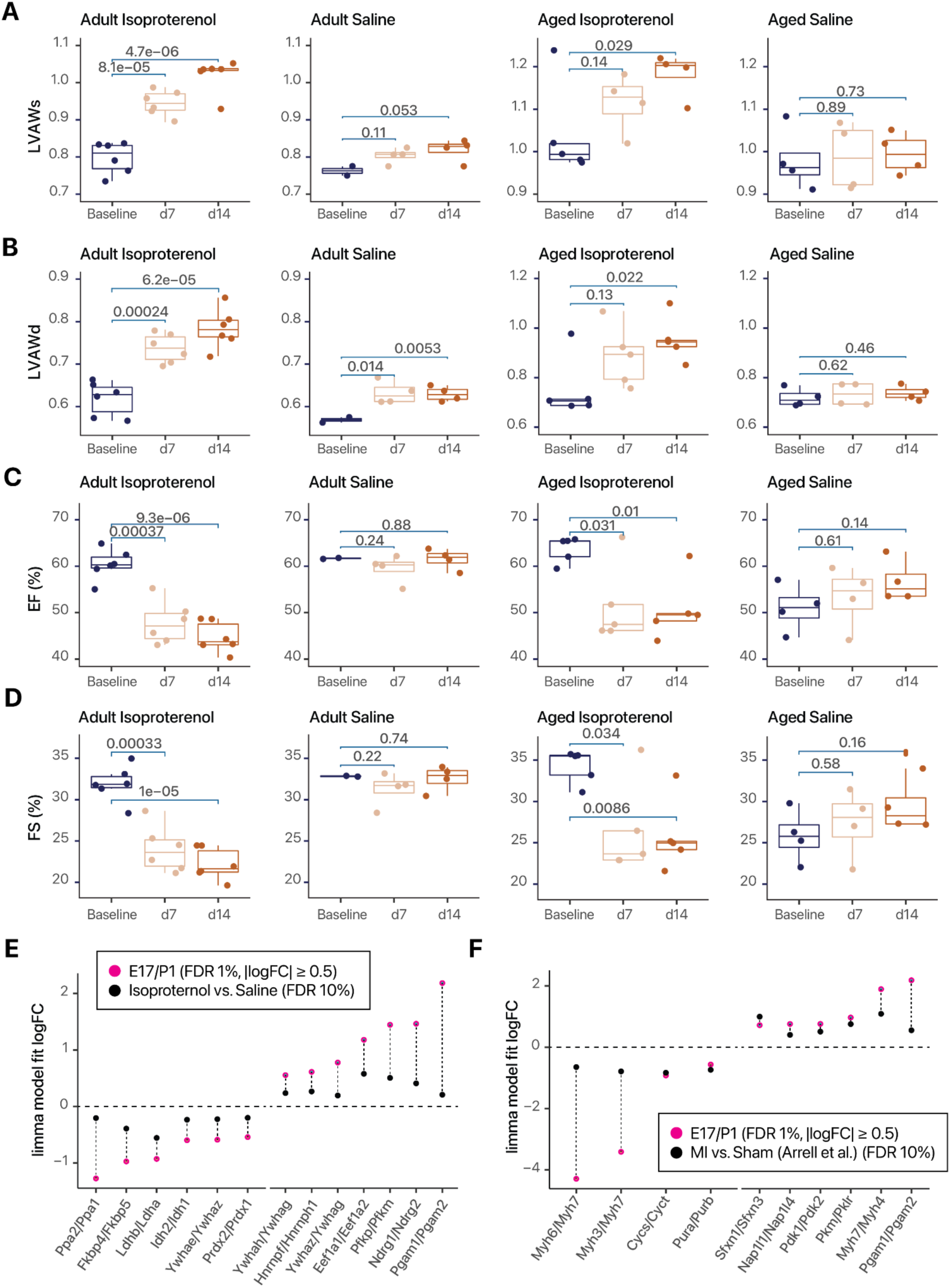
Partial return toward fetal isoform ratios in hypertrophic hearts. **A.** Echocardiographic measurements of left ventricular systolic anterior wall thickness (LVAWs) at baseline, day 7, and day 14 of isoproterenol or saline administration for young adult (left) and aged (right) C57BL/6 mice. P values obtained from two-sample t-tests are indicated above bars between respective comparisons of day 7 vs. baseline and day 14 vs. baseline. **B.** As in A, but for echocardiographic measurements of left ventricular anterior diastolic wall (LVAWd) thickness. **C.** As in A, but for ejection fraction (EF%) measurements. **D.** As in A, but for fractional shortening (FS%) measurements. **E.** Dotplot showing the limma fitted model logFC in fetal vs. perinatal heart comparisons (red) vs. adult hypertrophy vs. baseline hearts (black). Paralog pairs that are differentially expressed in fetal hearts (FDR 1%; abs(logFC) ≥ 0.5) and then showing significant co-directional changes in hypertrophic hearts (FDR 10%) are included. For hypertrophic hearts, n = 4–7 per age / treatment group. **F.** Dotplot showing limma fitted model logFC in fetal vs. perinatal heart comparisons (red) vs. mouse hearts 4 weeks after myocardial infarction (MI) vs. sham surgery in Arrell et al. Paralog pairs that are differentially expressed in fetal hearts (FDR 1%; abs(logFC) ≥ 0.5) and then showing significant co-directional changes in MI hearts (FDR 10%) are included. For MI hearts, n = 4 per group.

### Isoform ratio quantification in human cardiomyocyte differentiation

Finally, we asked whether the proteome-wide ratio test approach that we developed here can also be applied to other data sets as a generalizable method to gain new insights into gene regulation programs. To do so, we analyzed a set of TMT-labeled mass spectrometry data we previously generated to trace the proteomes of human induced pluripotent stem cells (hiPSC) under directed differentiation into cardiomyocytes (Lau et al., 2019). Briefly, three biological replicate lines of hiPSCs derived from different individual donors were directed to mesoderm specification, followed by cardiac progenitor and cardiomyocytes, using a small molecule-based GSK3β-inhibition Wnt-inhibition protocol (Lau et al., 2019). Protein profiles from the cells were then sampled daily from day 0 to day 14 of differentiation (**Table S10**). (data Retrieving the generated raw mass spectrometry data from PXD013426 and applying the paralog and splice isoform pair ratio calculation workflow described in this study, we then quantified the log ratios of 666 pairs of paralogs and 28 pairs of spliceoform proteins (**Table S11**). The resulting protein isoform ratios readily distinguished the cell samples by their differentiation days and stages in each line (**Figure 8A**). To quantify the isoform ratio shifts across major differentiation milestones, we then divided the samples into four groups: (1) hiPSC to mesoderm (day 0–2 of differentiation); (2) mesoderm to cardiac progenitor (day 3–6 of differentiation); (3) early cardiomyocyte (day 7–10 of differentiation); and (4) maturating cardiomyocyte (day 11–14 of differentiation). We then applied the linear model and empirical Bayes methods in limma to the generalized isoform log ratios as above (**Figure S6–S7; Table S12–S14**).

**Figure 8.**
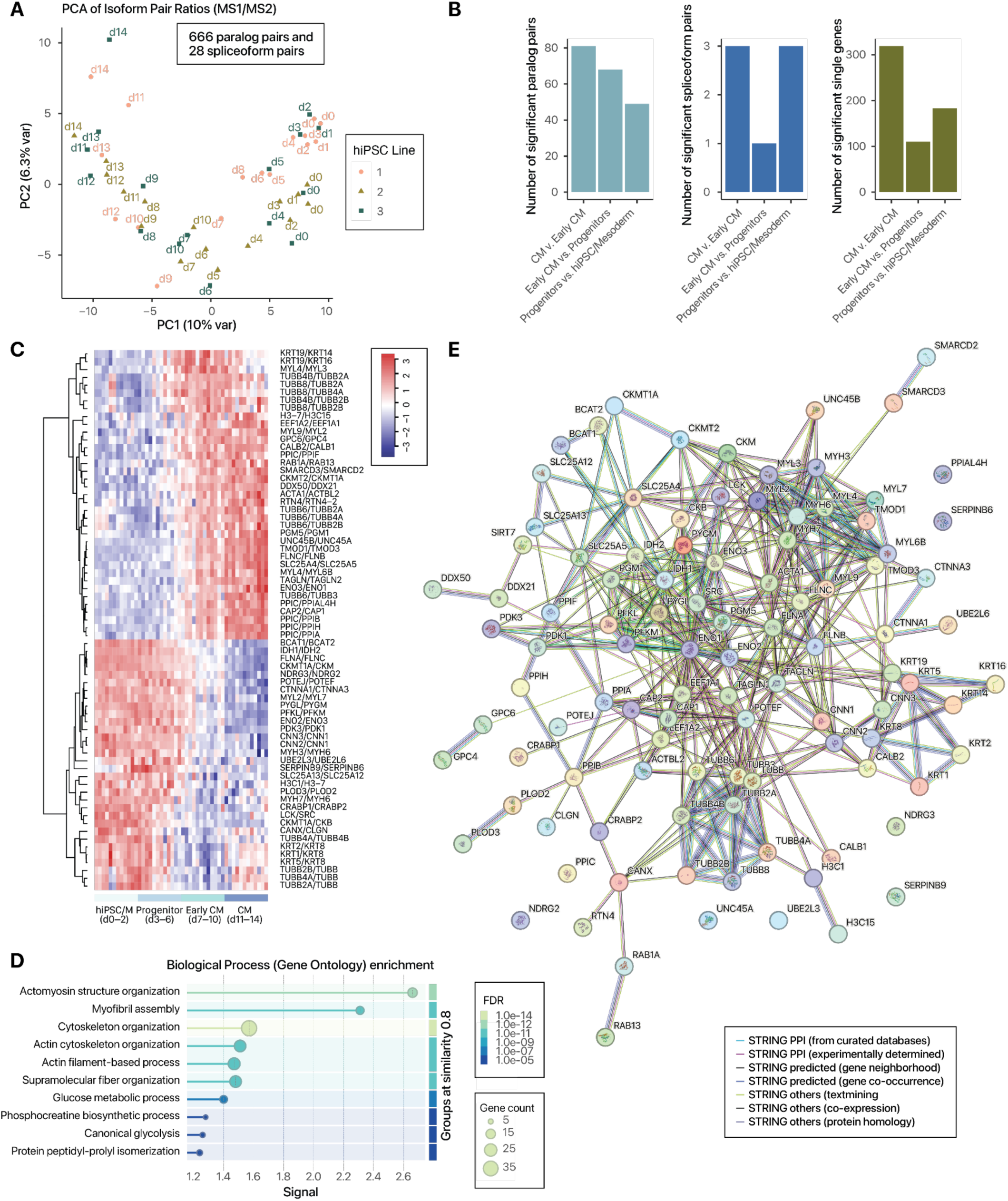
Proteome-wide protein isoform ratio quantification in hiPSC cardiomyocyte differentiation. **A.** Principal component analysis showing the ratios of 666 paralog pairs and 28 spliceoform pairs largely distinguished hiPSC differentiation stages. Colors: hiPSC biological replicate lines. **B.** Number of statistically significant (limma FDR adjusted P < 0.01; minor isoform fraction ≥ 0.05) paralog pairs (left); spliceoform pairs (middle); and individual proteins (right) at each stage of differentiation, from hiPSC/mesoderm (d0–2) to progenitors (d3–6), early cardiomyocyte (CM) (d7–10) and CM (d10–14). **C.** Heatmap showing isoform pairs with significantly different usage in early CM to CM transition. Colors: row standardized ratios. **D.** STRING enrichment graph of proteins involved in differential isoform usage. Colors: FDR. **E.** STRING network graph of proteins involved in differential isoform usage. Edge colors: STRING interaction type.

Notably, the results show that differentiation from progenitor cells to early cardiomyocytes appears to preferentially alter isoform usage over single-protein changes when contrasted with the changes in the other differentiation stages. This suggests that early cardiomyocyte specification represents a critical point of substantial isoform usage rewiring (**Figure 8B**). An inspection of the significant protein pairs at this stage reveals a number of isoform shifts that mirror mouse heart development, including differing usages in ENO2/ENO3, and MYH7/MYH6 (**Figure 8C**). On a pathway level, proteins with differential isoform usage in early cardiomyocytes are significantly enriched in actomyosin structure organization, myofibril assembly, glucose metabolic process, and related annotation terms (**Figure 8D**). This observation is corroborated by STRING network analysis, which show several distinct clusters of metabolic (e.g., PGM1, IDH1, PFKM), myofibril (MYL2, MYH6, MYH7), and cytoskeletal (e.g., tubulins, keratins, and calponins) proteins in interaction networks (**Figure 8E**). These results contrast with those from the mesoderm to progenitor stage, which instead features isoform usage changes in proteins associated with phosphocreatine process, as well as Wnt signaling pathway, but not cytoskeletal terms (**Figure S8**). On the other hand, early cardiomyocyte to cardiomyocyte transitions show a more prominent involvement of proteins in myofibril assembly and striated muscle development processes, but not metabolic terms (**Figure S9**). Taken together, we surmise that hiPSC cardiomyocyte differentiation is associated with several waves of isoform usage changes in each major milestone, each associated with different biological pathways and processes. Moreover, the results demonstrate that proteome ratio tests can provide insights into the proteome compositions of different biological systems.

## Discussion

This work presents to our knowledge the first proteomics study to systematically examine the ratiometric changes of protein isoform pairs in the fetal heart. To compare the absolute quantity of two proteins and thereby calculate their ratios across conditions, we have implemented a workflow that combines the MS1 precursor ion intensity of each peptide and the proportional abundance of their MS2 TMT reporter channels. This enables us to estimate the absolute abundance of individual proteins over 5 orders of magnitude. The results are highly consistent with other mass spectrometry experiments that used label-free quantification to derive absolute protein quantity in the mouse proteome, supporting that the MS1-MS2 deconvolution approach here is able to return consistently reliable protein quantification values.

The ratios between protein isoforms can reveal new complexities within biological systems, by discovering the reciprocally regulated relationships between two closely related homologous proteins. Paralog genes and alternative splicing present two major modalities from which new protein function is created. Paralogs are created from duplication events of ancestral genes or genomes that create redundant gene copies, which allow the ancestral copy to retain its conserved function while leaving the new copy free to take on new functional specialization under selection. These specializations may include the acquisition of differential regulation mechanisms by transcriptional and translational factors at different developmental stages, finetuning of dosage and expression specificity in different tissues, and the eventual divergence of molecular and physiological function (Mantica and Irimia, 2025). A deep parallelism has also been noted between the acquisition of new functions by differential gene regulation and by differential alternative splicing. Both mechanisms have harnessed duplication events (gene duplication and exon duplication) during bilateria evolution to allow extra copies of genetic material to evolve specialized function, and both can occur through new interactions between genetic elements (promoters and. splice sites) and protein regulators (transcription factors and. splice factors) in a context-specific manner (Mantica and Irimia, 2022). A protein (group) can also be regulated through the expression of both paralogs and alternative splicing isoforms (e.g., tropomyosins and troponins)

Applying the protein ratio tests to the fetal heart, we find numerous well established cardiac fetal gene pairs in the data. When measured in terms of the absolute magnitude of protein proportion swings, hallmark changes in contractile present the most prominent usage shifts in the fetal hearts (i.e., with the greatest log fold-changes), followed closely by glucose metabolism proteins. This may explain why previous work on the fetal heart program has placed a strong focus on these pathways. However, smaller magnitude changes can also effectuate biologically significant outcomes, and the ratios between two reciprocally regulated or mutually regulating proteins are sensitive to even relatively small differences in protein pool proportions, e.g., a mere 60/40 to 40/60 proportion shift would lead to an over 2-fold change in the relative ratio of the two proteins with respect to each other. Indeed, our results uncovered a large number of protein isoform pairs that to our knowledge have not been canonically established as part of the fetal gene program and that may help reveal new insights into cardiac development and disease. For instance, our data implicates a number of mitochondrial protein pairs in cardiac development, suggesting the shift away from carbohydrate metabolism as the preferred fuel source is paralleled by a concurrent remodeling of the mitochondrial protein composition. These changes involve paralog pairs such as inorganic pyrophosphatase PPA1/PPA2, ADP/ATP translocases ANT1/ANT2 (SLC25A4/SLC25A5), and VDAC1/VDAC2; as well as alternative isoform pairs like glutaminase-1 variants (KGA/GAC).

Changes in isoform usage could exert far reaching functional consequences by modulating protein enzymatic activity and interaction partners, even in cases where the isoforms share highly similar sequences and domain structures. For instance, while the β and α myosin heavy chains (MYH7 and MYH6) share a remarkable degree of sequence homology (McNally et al., 1989) (up to 93% identity in the mouse), the fast α isoform has up to threefold higher ATPase activity (Pope et al., 1980); the shift from MYH7 to MYH6 in postnatal heart is therefore consistent with adaptation toward higher contractile activity. In the case of enolase ENO1/ENO3 (∼84% sequence identity), enzymes purified from the liver (presumed ENO1 or alpha-/liver enolase) and the muscle (presumed ENO3 or beta-/muscle enolase) showed that under pH 7.4, both isozymes showed similar *K*_m_ with 2-phospho-D-glycerate in the forward glycolysis direction, but the muscle isozyme has a stronger affinity (lower *K*_m_) for phosphoenolpyruvate (Chang et al., 2021), which may suggest more versatile metabolic control in the postnatal heart. In the case of PCBP1 and PCBP2, two highly similar paralogs (∼85% sequence identity) that shift in proportion during fetal development, a domain-by-domain view suggests that the three RNA-binding KH domains (KH1 - residues 13–75 in PCBP1, KH2 - residues 97–162 in PCBP1, and KH3 - residues 279–343 in PCBP1) are highly conserved, whereas the linker between KH2 and KH3 are more variable. This suggests the core RNA-binding machinery of the proteins are nearly identical but the linker differences could affect flexibility between domains, or the ability to form interactions. Indeed, the over-expression of PCBP1 vs. PCBP2 in vitro led to highly correlated transcriptome profiles (r = 0.59, p ∼ 0) but also some differential effects, suggesting they affect similar but incompletely overlapping sets of genes.

Among mitochondrial proteins, the preference of ANT1 (SLC25A4) over SLC25A5 in the postnatal heart suggest adaptations of ATP export in a high metabolically active state, but may also have implications in cell death susceptibility given the implication of SLC25A4 in cardiac cell apoptosis (Zhou et al., 2024). Likewise, while the complex function and regulation of VDAC isoforms remains to be elucidated (Baines et al., 2007), there is evidence showing VDAC1 may promote apoptosis (Anflous et al., 2001) while VDAC2 may inhibit apoptosis (Cheng et al., 2003); hence the dual increase in VDAC1/VDAC2 and SLC25A4/SLC25A5 ratios could reflect a post-birth change in mitochondrial physiology that also alters cell death susceptibility. Further interpretation of the biological consequences of the isoform usage shifts would require functional studies. Such analysis may be aided by domain analysis to identify regions of sequence differences between isoforms; as well as consideration of isoform-specific enzyme activity or binding affinity, where such data is available.

In the hypertrophic mouse heart experiment, we find that only some fetal gene shifts are reversed and they do not revert fully to the protein pool compositions of E17 hearts when seen in the light of protein abundance proportions. Although the small sample size investigated here and the differences between animal disease models can both limit the number as well as magnitude of fetal gene reactivation that can be identified, the results nevertheless suggest that the return to the fetal gene program in pathological heart may be partial in amplitude at the protein level. This in turn may indicate there is considerable buffering of gene expression changes in post-transcriptional and post-translational layers. In a second application, the proteome-wide ratio tests in hiPSC cardiomyocyte differentiation revealed a number of shared changes in contractile and metabolic pairs between mouse heart development and various stages of human cardiac cell specification and maturation. This result suggests that a parallel shift in protein ratios occurs in human cells during cardiac differentiation processes while indicating the method would be broadly useful for the investigation of different species and models.

Finally, we consider some of the practical advantages that can come from explicitly testing for ratio differences rather than individual protein levels in proteomics experiments. In a recent work, Suhre suggested that in a genome association study, accounting for ratio associations with genetic variants can lead to significant increases in power (“p-gains”) by revealing the shared genetic and non-genetic variances between two genes, e.g., by uncovering a trans-QTL that simultaneously induces one gene while suppressing another (Suhre, 2024). In the case of quantitative proteomics, we posit that such a contradirectional modulator would have the effect of amplifying ratio changes over the effect size of individual protein fold-changes and by doing so increase the power of discovery. Moreover, in a mass spectrometry experiment, both latent biological factors (e.g., unaccounted for age or sex differences) as well as technical factors (e.g., chromatographic variations and batch effects) may exert a similar effect over two closely related proteins, which may then be effectively corrected by comparing their ratios between conditions. This self-normalization effect likely also motivated many historical biochemical studies to consider the ratios of two proteins in histological staining and immunoblots. Finally, by measuring the absolute abundance of isoforms within an isoform group, we reveal information about proportional pool sizes that can aid in data interpretation and prioritization. This contrasts with relying on the fold-change (i.e., up/down regulation) of individual proteins alone, which can obscure drastic differences in the copy numbers of two closely related proteins, in a manner analogous to the stoichiometric occupancy of post-translational modification sites vs. fold-change of modified peptides. In other words, a minor isoform may be expressed in such a miniscule level, that even up-regulated, may be of insufficient quantity to exert a functional outcome (e.g., a minor splice isoform that exists at 0.1% of the the canonical isoform will still only account for 0.4% of the protein pool after 4-fold up-regulation).

### Limitations of the Study

1. As we focus on comparing the ratios between two isoforms, this study omits isoforms that behave similarly to each other in fetal and adult hearts. Fetal genes that are not part of a paralog or spliceoform group (e.g., ANP) are likewise excluded.
2. MS2 TMT quantification can lead to ratio compression due to co-eluting isobaric peptides, which may bias measured ratios toward 1:1. This may be alleviated with the use of MS3 TMT quantification, but the benefits appear to be minor and minimal effects have been seen on the accuracy and interpretation of hyperplexed experiments (Klann et al., 2020; Savitski et al., 2018). The ratio comparison concept can be applied (and indeed more straightforwardly) to label-free quantification mass spectrometry data. Other general limitations of proteomics apply, including the still-incomplete coverage of all proteins, sample size constraints, and differences between bulk vs. single-cell proteomes. These issues can be addressed with advances in technology and instrumentation.
3. In the ratio tests, a protein measurement may appear in multiple ratios (i.e., Protein A over B and A over C) which introduces non-independence to the data matrix. To partially correct for this dependency, we used a more stringent 1% limma FDR adjusted P value, compared to the 5% threshold that is sometimes used in the literature. The effect of additional filtering and normalization steps is not investigated here.
4. Finally, additional experiments are needed to validate the novel fetal gene candidates. Isoform usage may also differ between humans and mice, e.g., α to β switch in human ventricles. While we showed using hiPSC data that the ratio tests can be applied to human cells, the translatability of the findings to human heart development remains to be elucidated. These limitations suggest valuable areas of future development.

## Conclusion

In summary, we describe a ratiometric proteomics workflow to discover differences in protein isoform usage across samples. Applying this workflow to the mouse heart, we recapitulate numerous hallmark fetal genes while discovering novel candidates, providing a resource of the fetal gene program of the heart at the protein level. More generally, the results highlight the prospects of applying protein ratio tests as a novel data analytical strategy of proteomics data, which should find utility in the discovery of gene regulatory mechanisms beyond individual-protein changes.

## Methods

### Animal Experiments

All animal protocols were approved by the Institutional Animal Care and Use Committee (IACUC) guidelines at the University of Colorado Denver/Anschutz Medical Campus. Fetal and postnatal C57BL/6J mouse hearts (n = 5 each) were acquired commercially at E17 and P1 from Zyagen (San Diego, CA), weighed, and homogenized for proteomics profiling (see below). For normal and hypertrophic adult mouse hearts, wild-type C57BL/6J mice were purchased from Jackson Laboratories (Bar Harbor, ME, USA) and housed in a temperature-controlled environment on a 12-h light/dark cycle and with free access to normal chow diet and water under National Institutes of Health (NIH) guidelines for the Care and Use of Laboratory Animals. Male adult (15-16 wk) and aged (83-84 wk) mice were administered with saline or 20 mg/kg isoproterenol by micro-osmotic pump (ALZET) implantation for a duration of 7 or 14 days as previously described (Lam et al., 2014) (n = 4, 7, 4, 6 for young adult saline 7 days, young adult isoproterenol 7 days, aged saline 7 days, aged isoproterenol 7 days; n = 4, 6, 4, 5 young adult saline 14 days, young adult isoproterenol 14 days, aged saline 14 days, aged isoproterenol 14 days, 40 male animals total). The animals were euthanized, followed by measurement of body weight, heart weight and tibia length. Left ventricles were collected and kept frozen at –80 °C.

#### Echocardiography

Transthoracic echocardiograms were performed using the VisualSonics Vevo2100 system (Visualsonics, Toronto, Canada). Animals were anesthetized in an induction chamber at 2% isoflurane. Once appropriately sedated, mice were transferred to the imaging platform. Hair on the chest was removed with a depilatory lotion and the mouse was secured by taping all four limbs to the platform. Body temperature was maintained at 37°C and isoflurane was delivered through a nose cone at 1.5% throughout the session. Mice were under anesthesia for no longer than 15 minutes. SAX views of the LV at the papillary muscle level were used to acquire M-mode images. Left ventricular anterior wall (LVAW) thickness in systole and diastole was measured using the M-mode images in order to measure changes in wall thickness and systolic function. Heart rate was measured in each image and averaged. All measurements were averaged from at least three cardiac cycles during the exhale phase and taken at body temperatures between 37°C and 38°C. All echocardiographic measurements and analyses were performed in a blinded manner.

#### Sex as a biological variable

For commercially procured fetal and perinatal hearts, specimens were not genotyped; it is unknown whether similar findings would be observed in male and female hearts. For the hypertrophy study, our study examined male mice to reduce potential variability in phenotype. It is unknown whether the findings are relevant for female mice.

### Protein Chemistry and Mass Spectrometry

Tissue homogenization was performed with a handheld homogenizer (OMNI International). Samples were placed on ice for 1 minute to allow cooling down. Repeat for a total of 3 rounds of homogenization. Homogenized tissues were sonicated with a handheld sonicator (Fisher Model 120), at 40% amplitude for 1 s, pause for 5 s, for 15 cycles. The samples were then centrifuged at 5,000 × g for 1 min at 4°C, and vortex for 10 s, and repeated for three rounds of sonication and centrifugation. The extracted protein samples were further centrifuged at 14,000 × g for 15 min at 4°C to collect supernatants and measure protein concentration with the BCA protein assay kit following manufacturer instructions. The samples were then reduced, alkylated, and digested with sequencing-grade trypsin as previously described (Han et al., 2022). The digested peptides were labeled with TMT 10-plex isobaric tags (Thermo), and fractionated using high-pH reversed-phase columns (Pierce) following manufacturer instructions.

Mass spectrometry was performed on a Q-Exactive HF orbitrap high-resolution mass spectrometer (Thermo) coupled to a Thermo Easy-nLC UPLC liquid chromatography system. For fetal heart samples, The LC gradient was as follows: 0 – 105 minutes, 0 – 40%B; 105 – 110 minutes, 40– 70% B, 110 – 115 minutes: 70 – 100% B; 115 – 120 minutes: hold; at 300 nL/min. Mass spectrometry data were acquired in data dependent acquisition (DDA) mode using typical settings including 60,000 MS1 and MS2 resolution, 3e6 MS1 AGC; 2e5 MS2 AGC; 100ms maximum IT; 15 TopN; isolation window 1.4 m/z; normalized CE 28, 30, 32; and 30 seconds dynamic exclusion. For cardiac remodeling samples, The LC gradient was as follows: 0 – 105 minutes, 0 – 30% B; 105 – 110 minutes, 30 – 70% B, 110 – 115 minutes: 70 – 100% B; 115 – 120 minutes: hold; at 300 nL/min. Mass spectrometry data were acquired in data dependent acquisition (DDA) mode using typical settings including 60,000 MS1 and MS2 resolution, 3e6 MS1 AGC; 2e5 MS2 AGC; 110 ms maximum IT; 15 TopN; isolation window 0.7 m/z; normalized CE 28, 30, 32; and 30 seconds dynamic exclusion.

### Data Analysis and Statistics

#### Mass spectrometry database search

Mass spectrometry data was searched using Sage v.0.14.5 (Lazear, 2023) aarch64-apple-darwin build to identify proteins and to measure peptide MS2 TMT and MS1 LFQ (label-free quantification) intensity, using typical settings including: missed_cleavages: 2, precursor_tol: –20 to +20 ppm; fragment_tol: –20 to +20 ppm; isotope: false; static_mods: C 57.0215, K 229.1629; variable mods: M 15.9949, Peptide N-terminus 229.1629, S 229.1629; generate_decoys: true; isotope_errors: 0 – 3; quant: tmt: Tmt10; quant: lfq: true; predict_rt: true. For mouse fetal and hypertrophy heart samples, the database used was retrieved on 2023-09-15 from UniProt (The UniProt Consortium et al., 2023) to contain 25,530 Swiss-Prot *Mus musculus* reviewed canonical and isoform entries. A peptide-level multiple testing adjusted q value cutoff of 0.01 was used for confident identification. For hiPSC data re-analysis, raw mass spectrometry files were retrieved from PXD013426. The database used was UniProt Swiss-Prot *Homo sapiens* canonical and isoform entries appended with JCAST as described in the original study (Lau et al., 2019). Sage settings include static_mods: C 57.0215, K 229.1629; variable mods: M 15.9949, Peptide N-terminus 229.1629; chimera: true; report_psms: 2. Other search settings were as described above. A peptide-level multiple testing adjusted q value cutoff of 0.01 was used for confident identification.

#### Calculation of MS1 and MS2 intensities

Peptides passing the q value cutoff and matching to non-decoy entries are used for quantification. For each peptide, their TMT reporter intensity is corrected for isotope spillover using non-negative least squares (Searle and Yergey, 2020) as described (Dostal et al., 2020), with the isotope purity matrix matching the TMT batch from the manufacturer. To overcome the issue of sequence redundancy, i.e., most peptides are not unique to a single database entry due to the high degree of similarities between canonical and alternative isoforms, we applied a protein isoform rollup routine for parsimony. The rationale of this workflow follows precedents in the protein inference literature (Savitski et al., 2015) and has also been employed in a prior study from our group (Currie et al., 2024). Briefly, it assumes that most alternative isoforms are not expressed or only present in very low concentration unless evidence suggests otherwise. Peptides that are shared across two proteins coded from the same gene are therefore assigned to the canonical form only, unless the alternative isoform possesses one or more uniquely mappable peptides in the database search result. All peptides that are uniquely mappable to a single UniProt isoform entry, or uniquely mappable to a single UniProt canonical entry and where the non-canonical entries have no unique peptides, are used for quantification. The TMT channel intensities of all constituent peptides for each protein for each sample are then summed.

We next retrieve the MS1 intensities of each peptide from the Sage “lfq.tsv” output, which are then filtered and summed for each protein as above. We then perform a first-pass protein MS1 quantity calculation. To account for a longer protein emitting give more label free intensity than an equimolar shorter protein, we divide the sum of MS1 intensities of all peptides that are uniquely mappable to a protein by the total protein sequence length retrieved from the FASTA database used in the search, which gives a near-identical normalization across log-log scale as division by number of theoretically observable tryptic peptides (data not shown).

Next, we calculate a second-pass distributed MS1 quantification value, by considering the intensities of all non-uniquely mapped peptides. Briefly, the MS1 intensities of peptides shared by multiple proteins are divided proportionally by the unique-peptide abundance as calculated n the first-pass (unique-peptide-only) MS1 quantification value of each of the shared protein, then added to the MS1 quantification value of each respective protein. The total MS1 intensity values of each sample are then normalized by column sum, and missing values are median-imputed. Finally, we distribute the MS1 quantification value of each protein across samples using the TMT matrix calculated above. The TMT matrix is row-wise normalized to 1 to compute a sample proportional abundance matrix, then multiplied to the second-pass MS1 quantification value to yield the final MS1-MS2 protein abundance. Rows (proteins) with only missing values (NA) are removed. For downstream statistical testing, the protein abundance values are normalized by variance stabilizing transformation to stabilize the mean-variance relationship in the data, by using the limma::normalizeVSN() function for the fetal data (which has only 1 TMT block). For the hypertrophy data, which has 5 TMT blocks and contains shared reference channels, an internal reference normalization using two reference channels is further carried out prior to variance stabilizing transformation. The normalized data are inspected using boxplots and PCA plots for all samples.

#### Comparison of protein abundance estimates

For comparison to other estimates of protein abundance, we convert the absolute quantification values to copy number per cells, based on 10^9^ total protein molecules per cell, and compared to to the iBAQ absolute quantification value-based copy number in NIH 3T3 mouse fibroblasts via identical gene names as reported in the Supplemental Table S3 of (Schwanhäusser et al., 2011). To further validate the MS1-MS2 based (i.e., TMT/LFQ) protein quantity values, we convert the composite protein quantities to proportional copy numbers/absolute abundances in p.p.m., and compared these values to absolute protein copy numbers in the mouse heart compiled in PaxDB (Huang et al., 2023), which is a protein quantity database compiled from the aggregation of existing data sets including mass spectrometry data from multiple studies in mouse heart tissues (Geiger et al., 2013; Huttlin et al., 2010; Lau et al., 2016).

#### Paralog and splice isoform identification

Paralogs are retrieved from the Ensembl (Martin et al., 2023) *Mus musculu*s data set using biomaRt (Durinck et al., 2009) with the getBM() function, retrieving the “ensembl_gene_id”, “external_gene_name”, “mmusculus_paralog_ensembl_gene”, “mmusculus_paralog_associated_gene_name”, “mmusculus_paralog_canonical_transcript_protein”, “mmusculus_paralog_orthology_type”, “mmusculus_paralog_subtype”, “mmusculus_paralog_perc_id” and “mmusculus_paralog_perc_id_r1” attributes. Using the “mmusculus_paralog_perc_id” and “mmusculus_paralog_perc_id_r1” fields, we then filter in only paralogs with 50% or higher bidirectional gene sequence identity between two paralogs within a pair. Paralog pairs that map to identical UniProt accessions or contain missing accessions are removed. Orthology type is retrieved from the “mmusculus_paralog_orthology_type” and “mmusculus_paralog_subtype” fields. We then identify all disjoint paralog groups within the paralog tables using gene names that are explicitly identified in the fetal proteomics experiment. Within each group, we then calculate the group size, average sequence identity, and the ratios of MS1-MS2 protein abundance within each sample.

The ratios of splice isoforms within isoform groups are calculated analogously, by identifying UniProt isoform accession (e.g., P#####-2, P#####-3, etc.) belonging to the same canonical UniProt accession (P#####). To corroborate the protein quantification values and to further identify splicing-derived isoforms, we further performed a database search using a computational strategy we previously developed to enable RNA-seq-guided proteomics to identify protein isoforms from the mass spectrometry data (Han et al., 2020; Lau et al., 2019). Briefly, a custom database was generated using JCAST v.0.3.4 (Ludwig and Lau, 2021) with deep RNA sequencing data generated from the mouse heart as described (Han et al., 2022). The non-canonical sequences were then appended to the Swiss-Prot canonical + isoform database to yield a total of 29,102 forward entries. This database was used to repeat the database search using MSFragger v.3.8 (Kong et al., 2017) using typical settings, including precursor_mass_lower: –20 ppm; precursor_mass_upper: 20 ppm; fragment_mass_tolerance 20 ppm; isotope error: 0/1/2; allowed_missed_cleavage: 2; num_enzyme_termini: 1. The search results then underwent confidence determination using MSBooster v.1.1.6 (Yang et al., 2023) and Percolator v.3.0 (The et al., 2016). The MS2 and MS1 intensity of confidently identified peptides (Percolator q value < 0.01) were then extracted using pyTMT v.0.4.1 (Dostal et al., 2020) and Riana v.0.7.1 (Hammond et al., 2022), respectively. Protein abundance for spliceoform pairs was then calculated as above.

#### Statistical tests and additional data analysis

For statistical testing, from the variance stabilization transformed data, the difference in normalized expression (x_1_ – x_2_) between two isoforms within a group were calculated to represent the generalized log ratio of the two proteins. The data table containing only either single-protein values or isoform ratios were then used for comparisons. To perform statistical tests of either single protein or isoform ratios, we used the empirical Bayes and moderated t-test implemented in the limma package v.3.58.1 (Ritchie et al., 2015) on the protein abundance data frame or the log difference of protein isoform data frame, respectively. The design matrix and linear model have a formula of expression ∼ age (E17 vs. P1) for the fetal/postnatal comparison; for hypertrophy: expression ∼ treatment (Isoproterenol vs. sham) + age (Young adult vs. old) + TMT block is used. Diagnostic plots for data linearity and distributions were performed using the built-in limma plotMA(), plotSA(), and plotMD() functions. Unless otherwise stated, an FDR adjusted limma adjusted P value cutoff of 0.01 (1% FDR) is considered significant for fetal comparisons, along with absolute log fold change (≥ 0.5). An FDR adjusted limma adjusted P of 0.1 (10% FDR) is considered significant for hypertrophy comparisons.

Functional enrichment analysis was performed using gene set enrichment analysis (GSEA) against Reactome annotated pathways (Gillespie et al., 2022) using the ReactomePA package (Yu and He, 2016). Enriched gene sets/annotations with FDR adjusted permutation test P value < 0.01 (1% FDR) are considered significant. Additional data analysis is performed with the aid of common R packages in the tidyverse and on Bioconductor, and the STRING-db (Szklarczyk et al., 2023, p. 2) website functions.

### Immunoblots

Hearts from E17 and P1 stage C57BL/6 mice (n=3 each) were lysed in RIPA buffer (Cat# 89901, Thermo Scientific) supplemented with protease inhibitor (Cat# 78442, Thermo Scientific) and homogenized using a bead mill homogenizer (Omni Bead Ruptor 4) at speed 5 for 20 seconds. Homogenized tissues were then further sonicated using a handheld sonicator probe for 15 pulses (1s on/1s off, 40% amplitude). Crude lysates were centrifuged at 13, 500 g for 15 minutes and supernatant was collected into a clean tube. Digital immunoblot analysis of the protein lysates was performed in the Jess Simple Western System (ProteinSimple) using the 12–230 kDa Separation module (Cat# SM-W004, ProteinSimple), according to the manufacturer’s instructions. The following primary antibodies and the dilution ratios were used in the Jess Simple Western: ENO1 (Cat# 11204-1-AP, Proteintech; 1:100), ENO3 (Cat# 55234-1-AP, Proteintech; 1:200), PGAM1 (Cat# NBP1-49532, Novus Biologicals; 1:40), PGAM2 (Cat# 15550-1-AP, Proteintech; 1:10), PPA-1 (Cat# 14985-1-AP, Proteintech; 1:10), PPA-2 (Cat# 16662-1-AP, Proteintech; 1:50), GAC-specific (Cat# 19958-1-AP, Proteintech; 1:50), and KGA-Specific antibody (Cat# 20170-1-AP, Proteintech; 1:50). Normalization was performed using a protein normalization reagent (Cat# DM-PN02, ProteinSimple) according to the manufacturer’s guidelines. Data analysis was performed using the Compass software (version 6.1.0) for Simple Western. Unless otherwise stated, a two-sample, two-tailed t-test is used for statistical tests for immunoblots data.

### RNA sequencing

AC16 cells were seeded into 6-well plates in DMEM/F12 supplemented with 10% fetal bovine serum. Next day, when the cells reached ∼70% confluency, they were transfected with PCBP1::Halo plasmid, PCBP2::Halo plasmid, or both at a concentration of 1 mg of each plasmid in 2 mL media using Lipofectamine 2000 (Cat# 11668019, Invitrogen) transfection reagent. Untransfected cells served as control. After 6 hours of incubation, the media containing the transfection reagent was replaced with fresh media. Cells were lysed with Trizol (Cat# 15596018, Invitrogen) 48 hours post-transfection. Lysates were centrifuged at 13500 × g for 5 minutes and RNA extraction was performed using Direct-zol RNA Kit (Cat# R2050, Zymo). RNA yield was measured using a Qubit fluorometer (Thermo) and RNA integrity score was measured using TapeStation (Agilent). PolyA enrichment, library construction, and short-read sequencing were carried out at Novogene USA. Sequencing was performed in paired-end mode with a read length of 150 base pairs and depth of ∼20 G per sample on an Illumina NovaSeq platform. The RNA-seq data were then aligned against GRCh38.p14/GENCODE v47 (Frankish et al., 2021) using STAR v.2.7.11a (Dobin et al., 2013), with –sjdbOverhang 149 and typical settings. Transcript assembly was performed using StringTie v.2.1.1 (Pertea et al., 2015), and differential expression analysis using DESeq2 (Love et al., 2014) using typical settings. Results with apeglm (Zhu et al., 2019) shrinkage s-value < 0.05 and abs(logFC) > 0.25 are considered significant.

## Author Contributions

Y.H., S.B., S.A.W., and M.P.L. designed the original investigations and acquired the data. E.L. and M.P.L analyzed the data and drafted the manuscript. B.P. and D.N. contributed additional research designs. All authors contributed to the submitted version of the manuscript. E.L. and M.P.L. handled funding.

## Acknowledgments

This work was supported in part by NIH awards R35GM146815 and R00HL144829 to E.L; R01HL169473 to E.L and M.P.L; R01GM144456, and R01HL141278 to M.P.L; AHA Postdoctoral

Fellowship 25POST1372064 to B.P.

## Supplemental Data

**Table S1** - Data of all quantified absolute protein abundance in the mouse E17 vs. P1 heart data set

**Table S2** - Data of all quantified protein isoform ratios in the mouse E17 vs. P1 heart data set

**Table S3** - Results of statistical tests of protein paralog ratios in the mouse E17 vs. P1 heart data set

**Table S4** - Results of statistical tests of individual protein expression in the mouse E17 vs. P1 heart data set

**Table S5** - Protein paralog pairs with significant shifts in isoform usage but not individual protein expression

**Table S6** - Evolutionary conservation and sequence identity of Ensembl retrieved paralogs from all quantified paralog groups

**Table S7** - Alternative splicing isoforms with significant difference in in the mouse E17 vs. P1 heart data set

**Table S8** - Results of statistical tests of individual protein expression in the mouse isoproterenol-induced hypertrophy data set

**Table S9** - Results of statistical tests of protein isoform ratios in the mouse isoproterenol-induced hypertrophy data set

**Table S10** - Data of all quantified absolute protein abundance in the hiPSC differentiation data set

**Table S11** - Data of all quantified protein isoform ratios in the hiPSC differentiation data set

**Table S12** - Results of statistical tests of protein isoform ratios in hiPSC/mesoderm vs. cardiac progenitor cells

**Table S13** - Results of statistical tests of protein isoform ratios in early hiPSC-cardiomyocytes vs. cardiac progenitor cells

**Table S14** - Results of statistical tests of protein isoform ratios in hiPSC-cardiomyocytes vs. early hiPSC-cardiomyocytes

## Supplemental Figures

**Figure S1.**
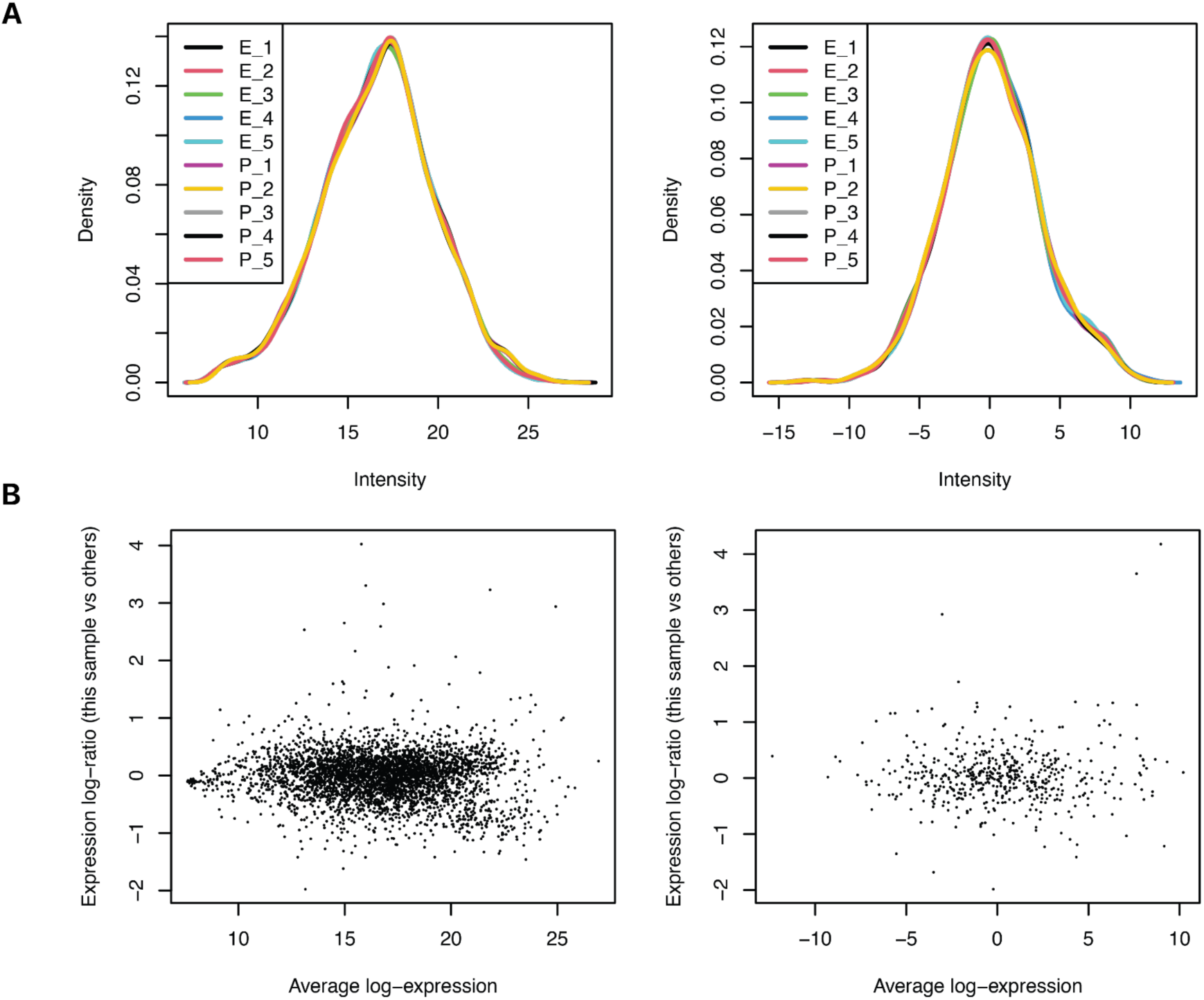
Comparison of limma diagnostic plots in protein and ratio analysis. **A.** Distribution of generalized log intensity among all individual protein species (left) and protein paralog ratios (right). **B.** Average log-expression vs. log-ratio of a representative sample (E17 Rep 4) against all other samples among all individual protein species (left) and protein paralog ratios (right) in the E17 (E) vs. P1 (P) mouse heart experiments.

**Figure S2.**
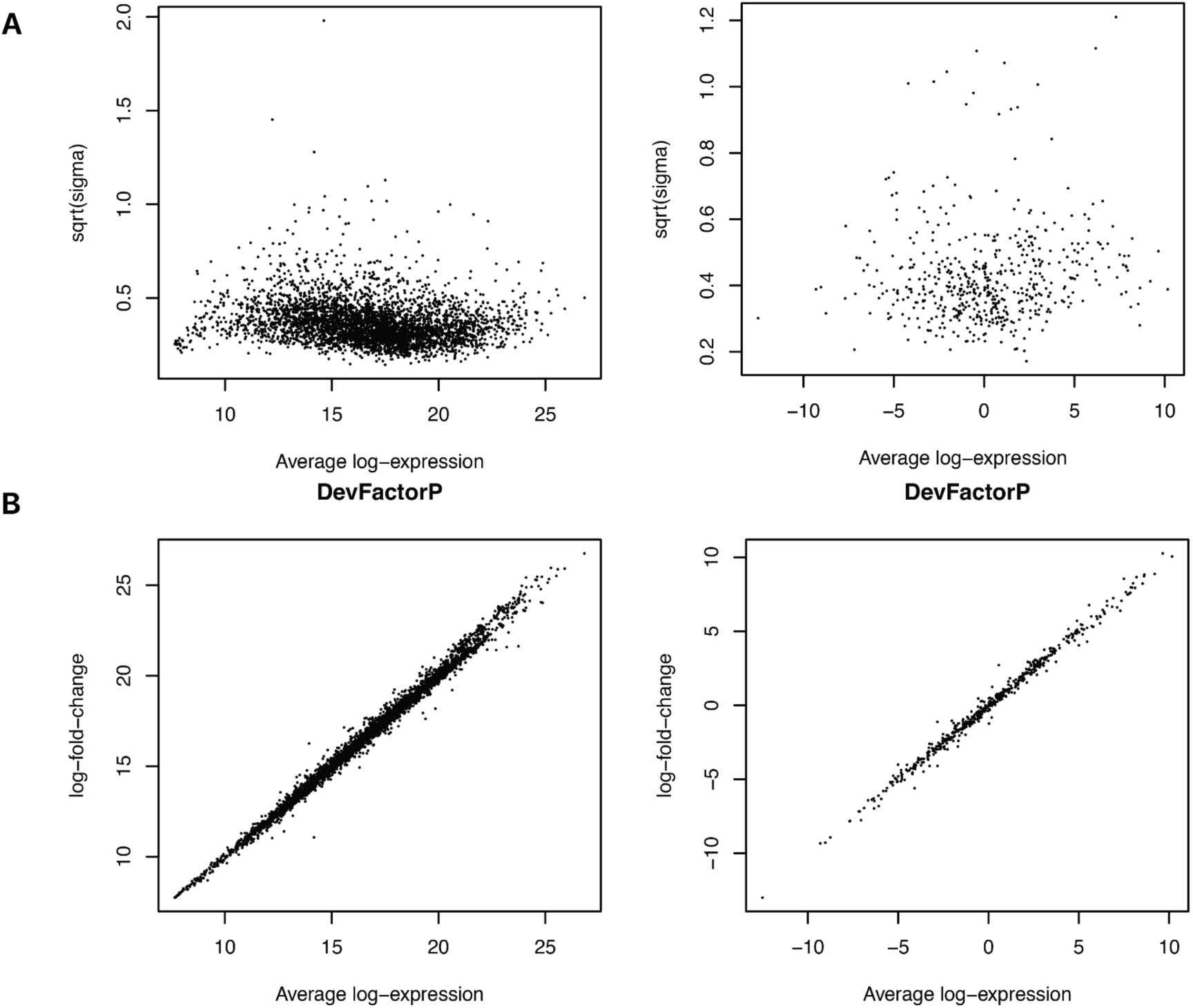
Comparison of limma diagnostic plots in individual protein species and ratio analysis (continued). **A.** Average log-expression vs. fitting residual variance (Sigma) after linear model fitting, among all individual protein species (left) and protein paralog ratios (right) in the E17 (E) vs. P1 (P) mouse heart experiments. **B.** Log expression over fold-change plot for developmental stage factor (perinatal) between all individual protein species (left) and protein paralog ratios (right).

**Figure S3.**
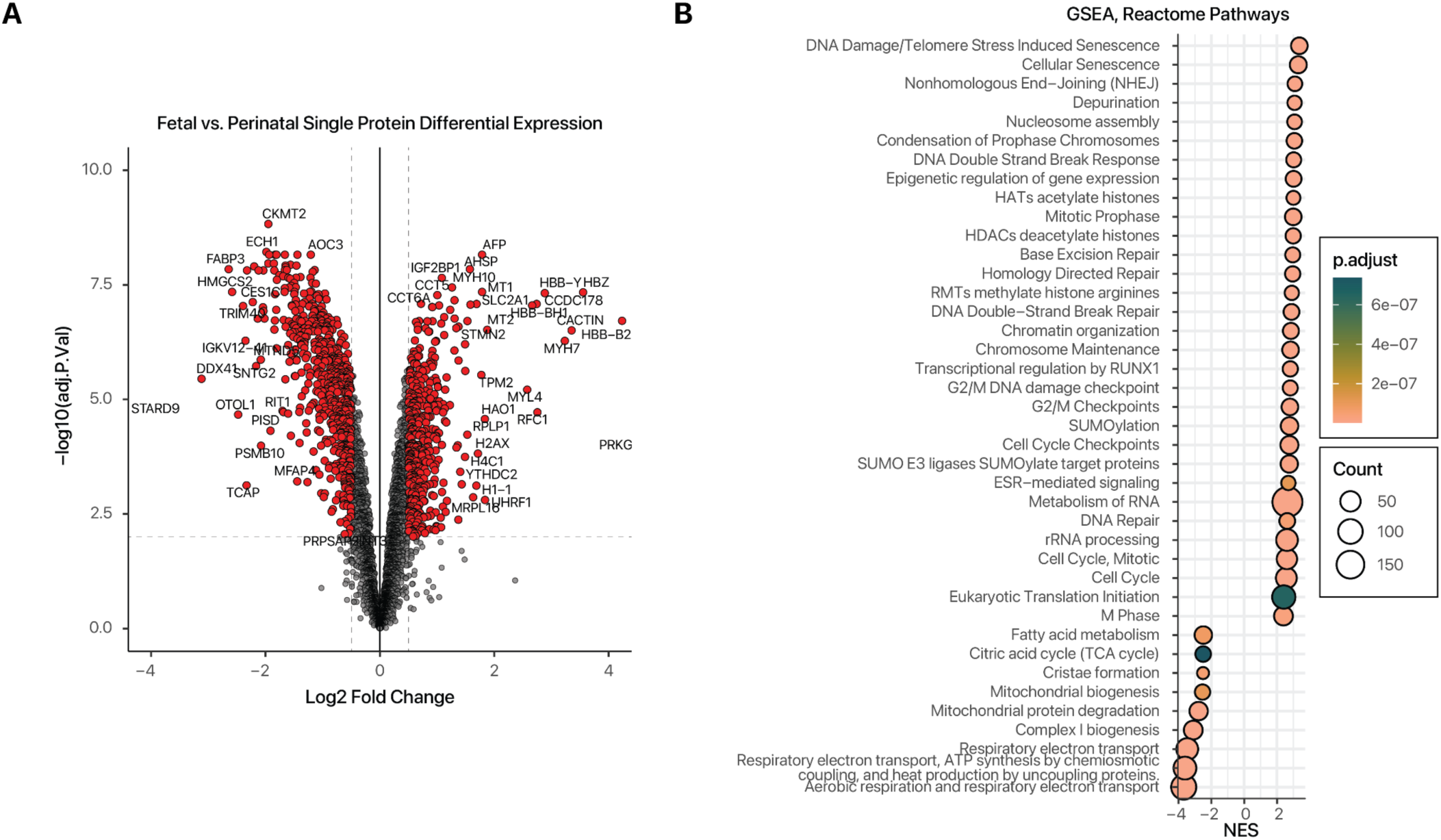
Individual protein abundance changes in fetal vs. perinatal hearts. **A.** Volcano plot showing the log2 fold-change (x-axis) vs. –log10 P value (y-axis) of compared individual proteins in E17 vs. P1 hearts. **B.** Gene set enrichment analysis (GSEA) of protein fold-change (E17/P1) against Reactome pathways. Color: GSEA adjusted P values; x-axis: GSEA normalized enrichment score (NES); size: number of proteins mapped to pathway.

**Figure S4.**
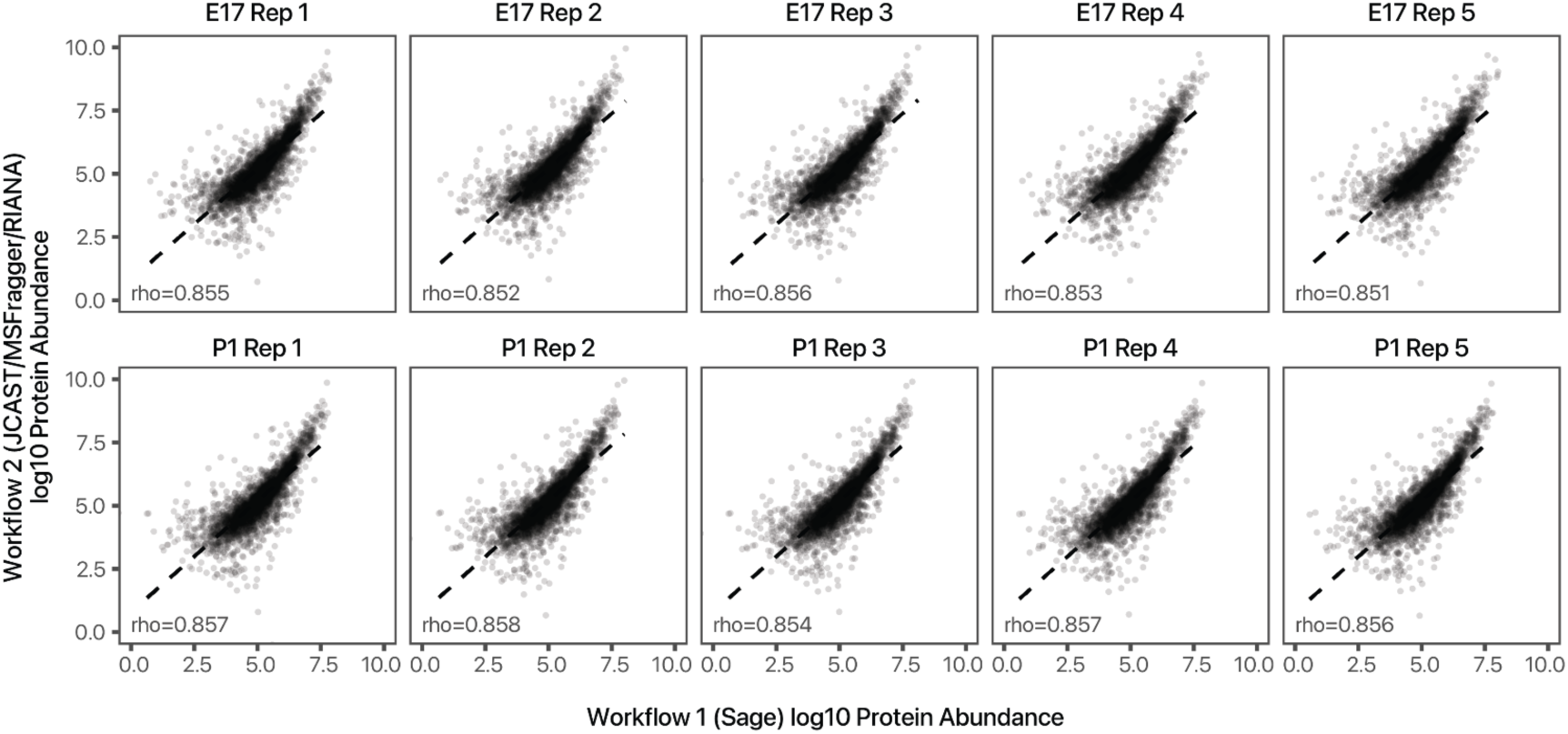
Comparison of protein quantification workflows. Scatter plots showing the measured protein abundance (MS1 and MS2 combined) in (x-axis) the Sage/Swiss-Prot workflow and (y-axis) the JCAST/MSFragger workflow. Robust correlation (Spearman’s correlation coefficient rho: ∼0.85) was observed across ten samples over 5 orders of magnitude in protein abundance in the fetal/postnatal comparisons between the two workflows, despite the use of different search engine, post-processing step, MS1 peak area integration, and TMT intensity extraction tools.

**Figure S5.**
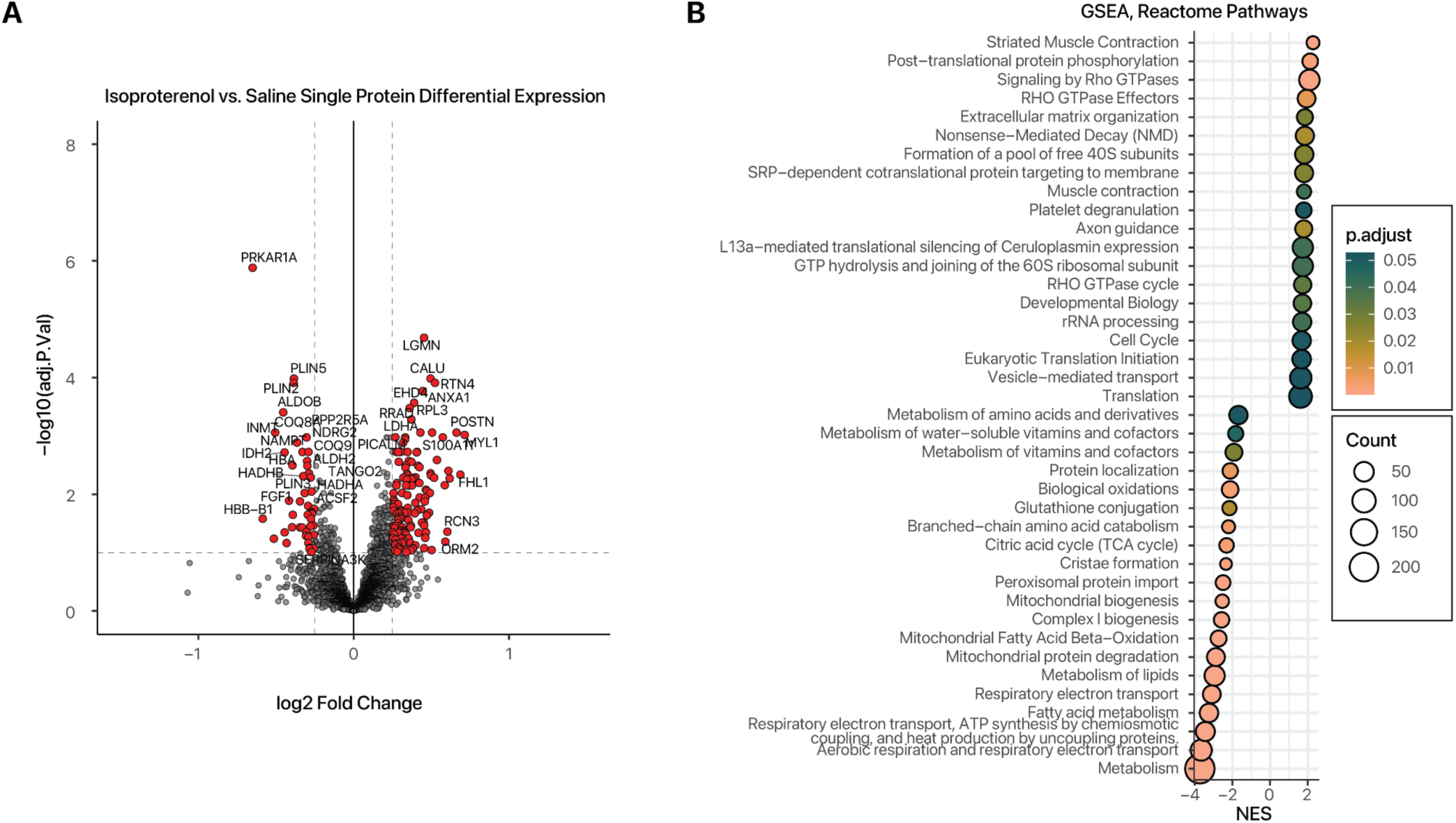
Individual protein abundance changes in hypertrophy vs. normal hearts. **A.** Volcano plot showing the log2 fold-change (x-axis) vs. –log10 P value (y-axis) of compared individual proteins (hypertrophy/normal). **B.** Gene set enrichment analysis (GSEA) of protein fold-change (hypertrophy/normal) against Reactome pathways. Color: GSEA adjusted P values; x-axis: GSEA normalized enrichment score (NES); size: number of proteins mapped to pathway.

**Figure S6.**
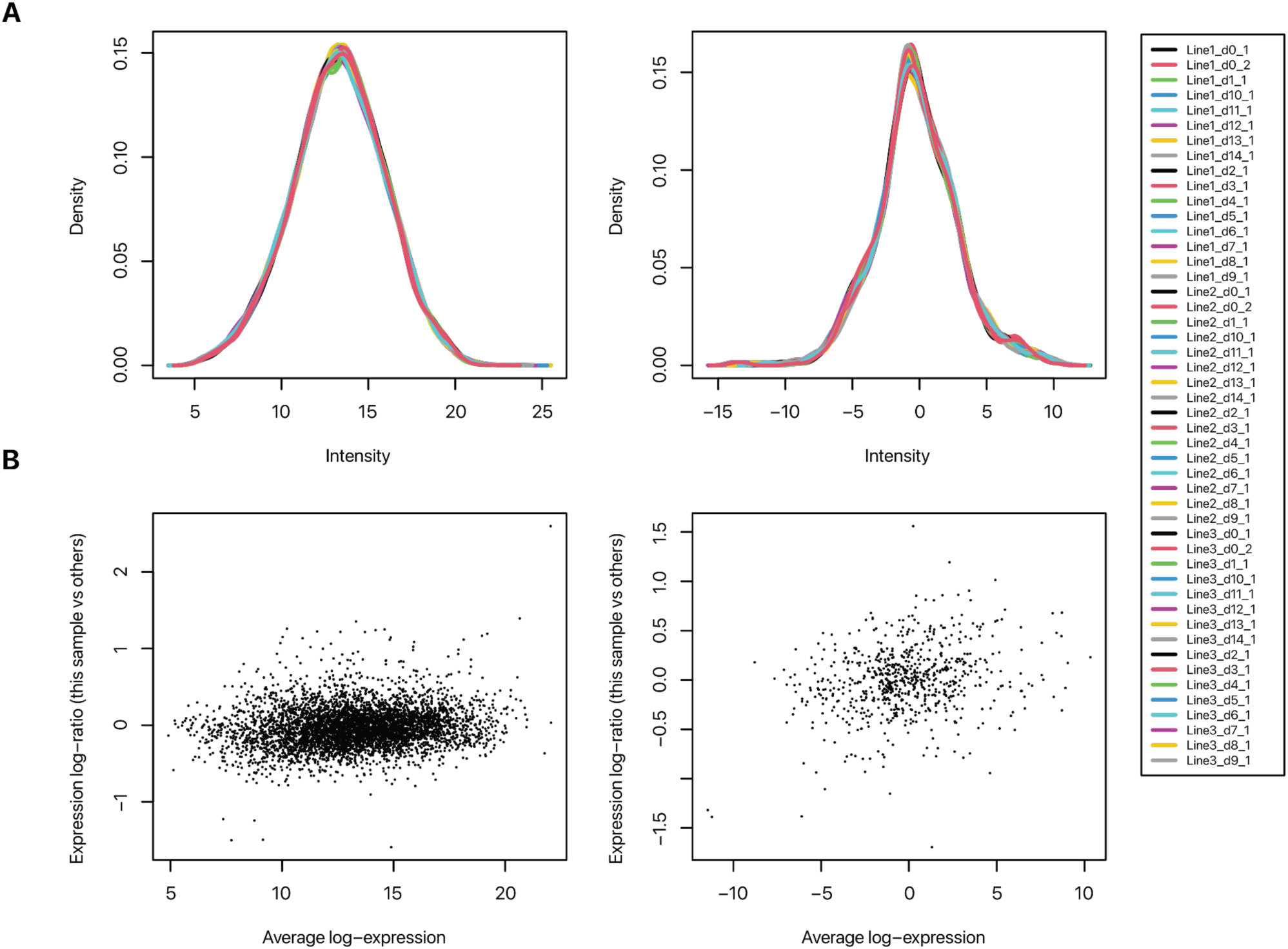
hiPSC limma diagnostic plots. **A.** Distribution of generalized log intensity among all individual protein species (left) and protein paralog ratios (right). **B.** Average log-expression vs. log-ratio of a representative sample (Line1_d10_1) against all other samples among all individual protein species (left) and protein paralog ratios (right) in the hiPSC differentiation data set.

**Figure S7.**
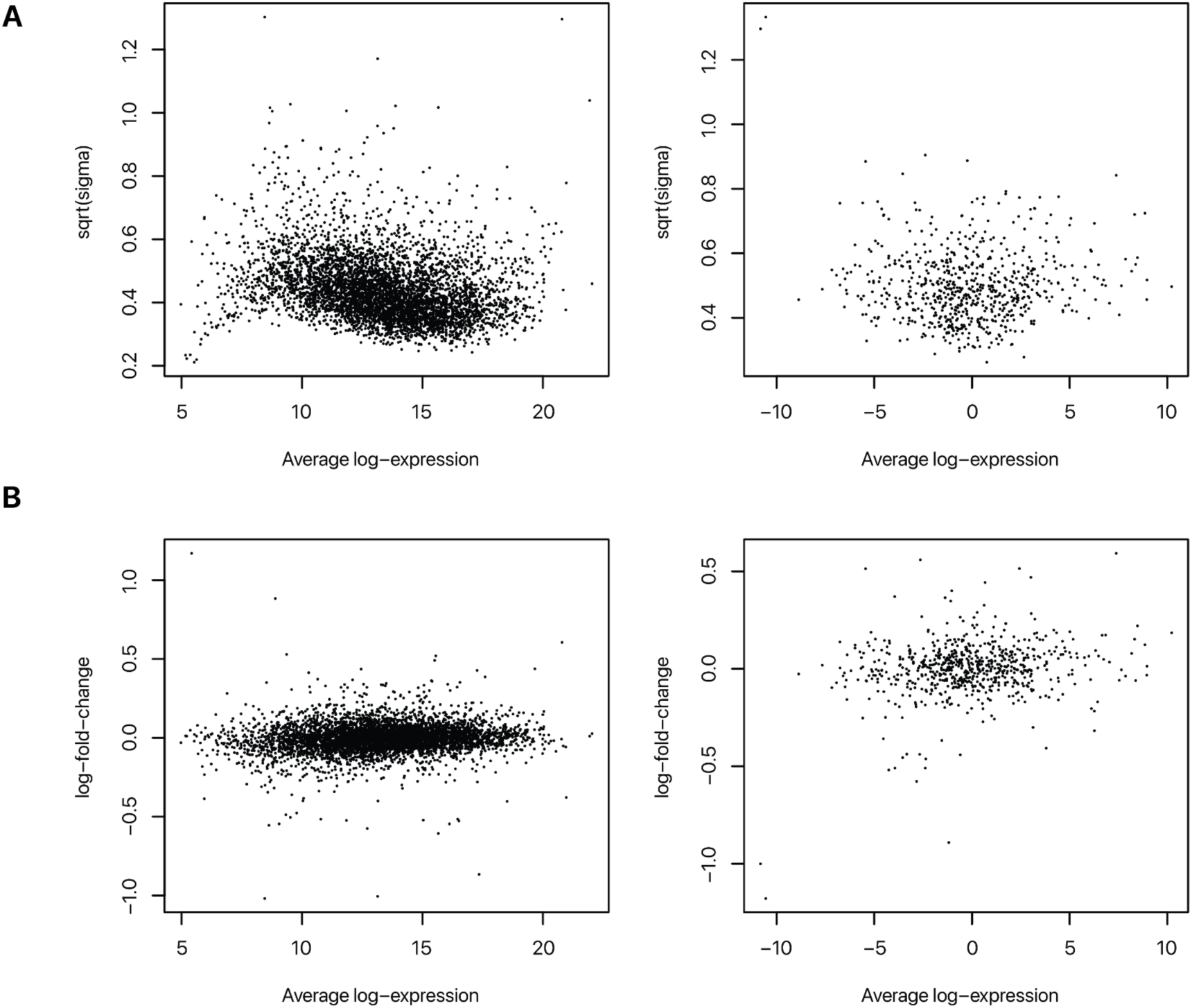
hiPSC limma diagnostic plots (continued). **A.** Average log-expression vs. fitting residual variance (Sigma) after linear model fitting, among all individual protein species (left) and protein paralog ratios (right) in the hiPSC differentiation data set. **B.** Log expression over fold-change plot for line factor (line 3) between all individual protein species (left) and protein paralog ratios (right).

**Figure S8.**
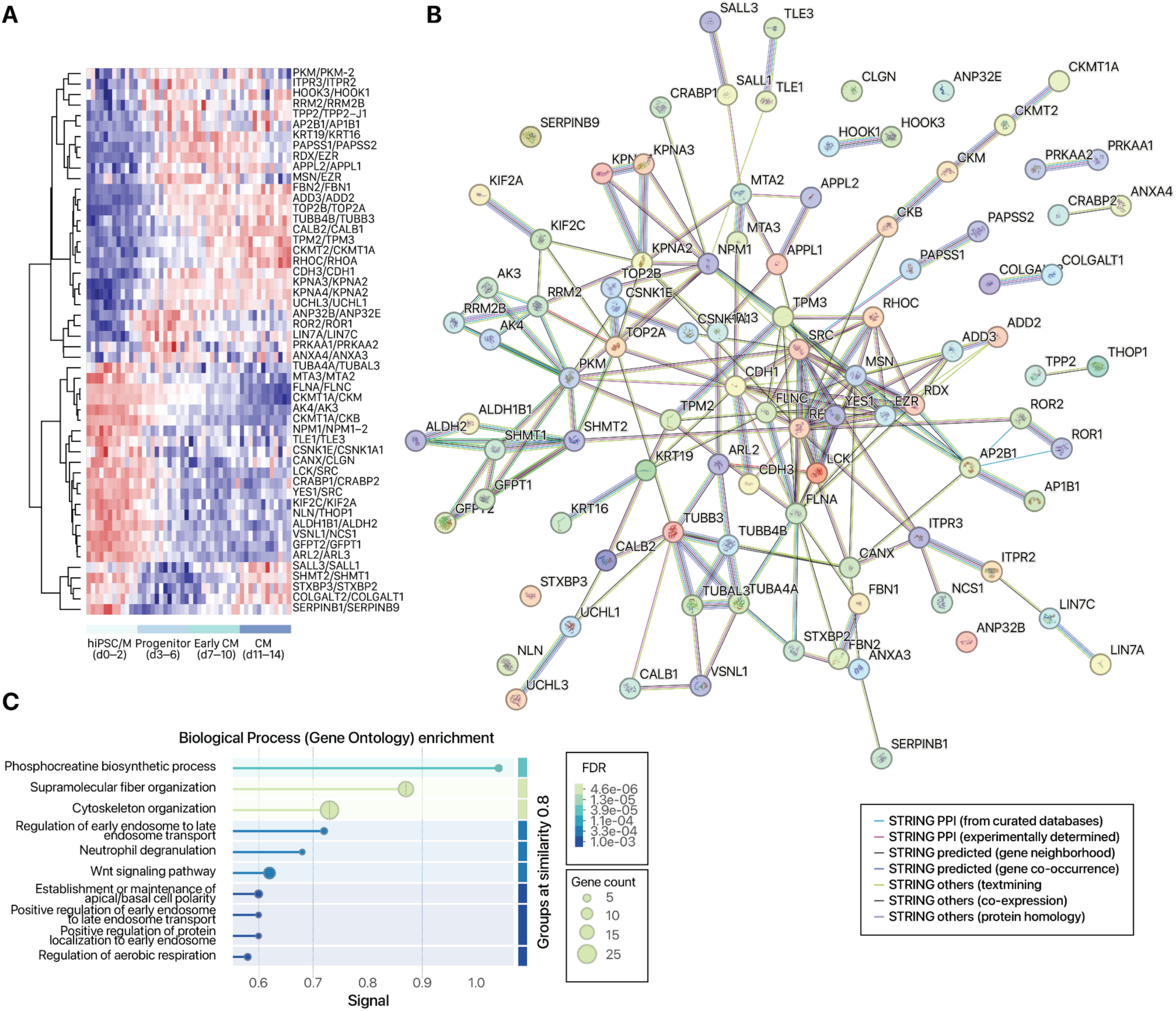
Differential protein isoform usage in hiPSC/mesoderm to cardiac progenitor transition. **A.** Heatmap showing isoform pairs with significantly different usage in hiPSC/mesoderm to progenitor transition. Colors: row standardized ratios. **B.** STRING network graph of proteins involved in differential isoform usage. Edge colors: STRING interaction type. **C.** STRING enrichment graph of proteins involved in differential isoform usage. Colors: FDR.

**Figure S9.**
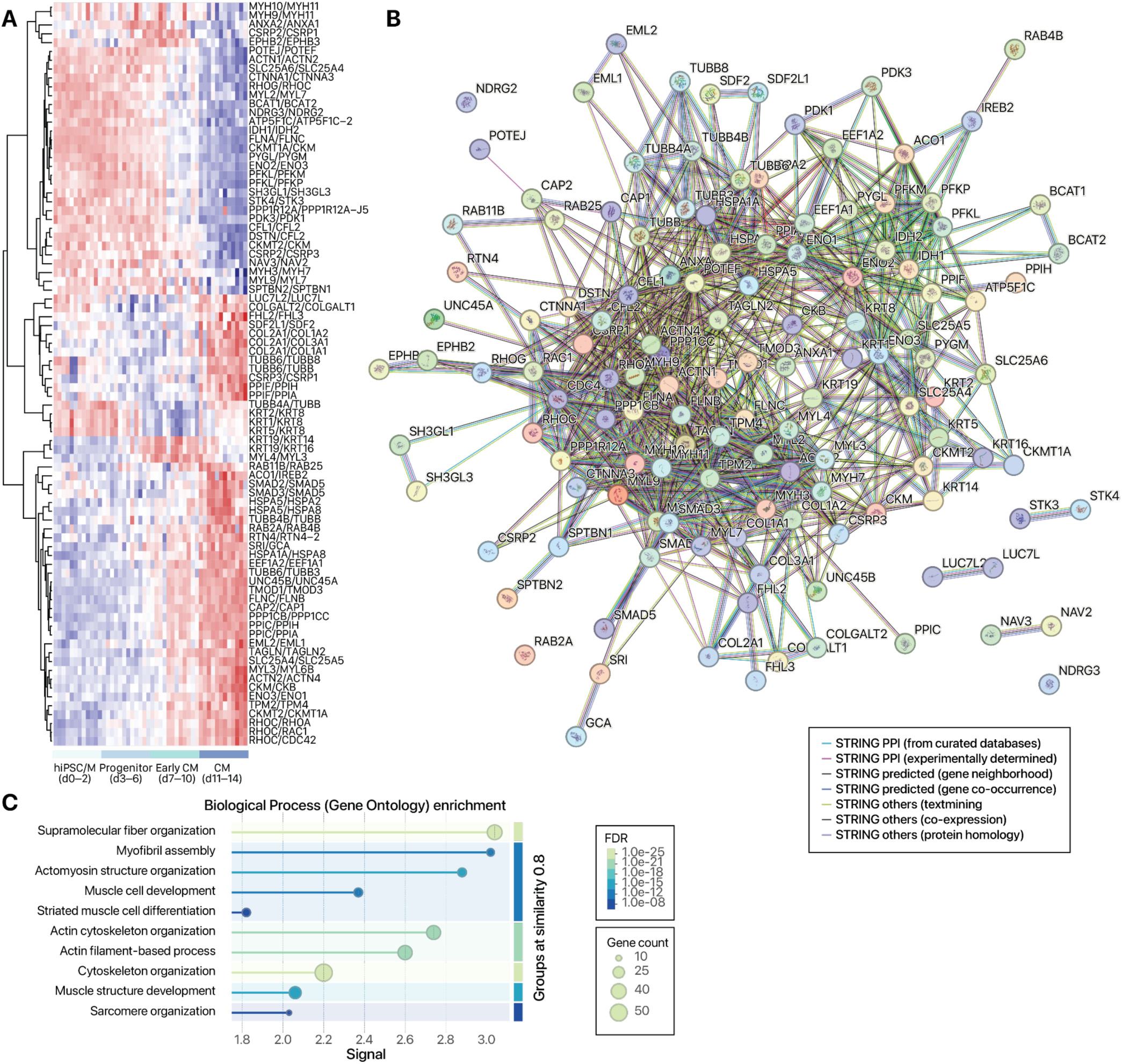
Differential protein isoform usage in early CM to CM transition. **A.** Heatmap showing isoform pairs with significantly different usage in early CM to CM transition. Colors: row standardized ratios. **B.** STRING network graph of proteins involved in differential isoform usage. Edge colors: STRING interaction type. **C.** STRING enrichment graph of proteins involved in differential isoform usage. Colors: FDR.

## References

Acoba, M.G., Alpergin, E.S.S., Renuse, S., Fernández-del-Río, L., Lu, Y.-W., Khalimonchuk, O., Clarke, C.F., Pandey, A., Wolfgang, M.J., Claypool, S.M., 2021. The mitochondrial carrier SFXN1 is critical for complex III integrity and cellular metabolism. Cell Rep. 34, 108869. 10.1016/j.celrep.2021.108869

Adamoski, D., M. Dos Reis, L., Mafra, A.C.P., Corrêa-da-Silva, F., Moraes-Vieira, P.M.M.D., Berindan-Neagoe, I., Calin, G.A., Dias, S.M.G., 2024. HuR controls glutaminase RNA metabolism. Nat. Commun. 15, 5620. 10.1038/s41467-024-49874-x

Ahrné, E., Martinez-Segura, A., Syed, A.P., Vina-Vilaseca, A., Gruber, A.J., Marguerat, S., Schmidt, A., 2015. Exploiting the multiplexing capabilities of tandem mass tags for high-throughput estimation of cellular protein abundances by mass spectrometry. Methods San Diego Calif 85, 100–107. 10.1016/j.ymeth.2015.04.032

Anflous, K., Armstrong, D.D., Craigen, W.J., 2001. Altered Mitochondrial Sensitivity for ADP and Maintenance of Creatine-stimulated Respiration in Oxidative Striated Muscles from VDAC1-deficient Mice. J. Biol. Chem. 276, 1954–1960. 10.1074/jbc.M006587200

Apt, D., Watts, R.M., Suske, G., Bernard, H.U., 1996. High Sp1/Sp3 ratios in epithelial cells during epithelial differentiation and cellular transformation correlate with the activation of the HPV-16 promoter. Virology 224, 281–291. 10.1006/viro.1996.0530

Arrell, D.K., Rosenow, C.S., Yamada, S., Behfar, A., Terzic, A., 2020. Cardiopoietic stem cell therapy restores infarction-altered cardiac proteome. NPJ Regen. Med. 5, 5. 10.1038/s41536-020-0091-6

Baines, C.P., Kaiser, R.A., Sheiko, T., Craigen, W.J., Molkentin, J.D., 2007. Voltage-dependent anion channels are dispensable for mitochondrial-dependent cell death. Nat. Cell Biol. 9, 550–555. 10.1038/ncb1575

Barbosa-Morais, N.L., Irimia, M., Pan, Q., Xiong, H.Y., Gueroussov, S., Lee, L.J., Slobodeniuc, V., Kutter, C., Watt, S., Çolak, R., Kim, T., Misquitta-Ali, C.M., Wilson, M.D., Kim, P.M., Odom, D.T., Frey, B.J., Blencowe, B.J., 2012. The Evolutionary Landscape of Alternative Splicing in Vertebrate Species. Science 338, 1587–1593. 10.1126/science.1230612

Belkin, A.M., Zhidkova, N.I., Balzac, F., Altruda, F., Tomatis, D., Maier, A., Tarone, G., Koteliansky, V.E., Burridge, K., 1996. Beta 1D integrin displaces the beta 1A isoform in striated muscles: localization at junctional structures and signaling potential in nonmuscle cells. J. Cell Biol. 132, 211–226. 10.1083/jcb.132.1.211

Bergeron, S.E., Zhu, M., Thiem, S.M., Friderici, K.H., Rubenstein, P.A., 2010. Ion-dependent polymerization differences between mammalian beta- and gamma-nonmuscle actin isoforms. J. Biol. Chem. 285, 16087–16095. 10.1074/jbc.M110.110130

Bhavsar, P.K., Dhoot, G.K., Cumming, D.V., Butler-Browne, G.S., Yacoub, M.H., Barton, P.J., 1991. Developmental expression of troponin I isoforms in fetal human heart. FEBS Lett. 292, 5–8. 10.1016/0014-5793(91)80820-s

Blencowe, B.J., 2017. The Relationship between Alternative Splicing and Proteomic Complexity. Trends Biochem. Sci. 42, 407–408. 10.1016/j.tibs.2017.04.001

Castelló, A., Rodríguez-Manzaneque, J.C., Camps, M., Pérez-Castillo, A., Testar, X., Palacín, M., Santos, A., Zorzano, A., 1994. Perinatal hypothyroidism impairs the normal transition of GLUT4 and GLUT1 glucose transporters from fetal to neonatal levels in heart and brown adipose tissue. Evidence for tissue-specific regulation of GLUT4 expression by thyroid hormone. J. Biol. Chem. 269, 5905–5912. 10.1016/S0021-9258(17)37547-6

Chang, A., Jeske, L., Ulbrich, S., Hofmann, J., Koblitz, J., Schomburg, I., Neumann-Schaal, M., Jahn, D., Schomburg, D., 2021. BRENDA, the ELIXIR core data resource in 2021: new developments and updates. Nucleic Acids Res. 49, D498–D508. 10.1093/nar/gkaa1025

Chen, X., Liu, Ying, Xu, C., Ba, L., Liu, Z., Li, Xiuya, Huang, J., Simpson, E., Gao, H., Cao, D., Sheng, W., Qi, H., Ji, H., Sanderson, M., Cai, C.-L., Li, Xiaohui, Yang, L., Na, J., Yamamura, K., Liu, Yunlong, Huang, G., Shou, W., Sun, N., 2021. QKI is a critical pre-mRNA alternative splicing regulator of cardiac myofibrillogenesis and contractile function. Nat. Commun. 12, 89. 10.1038/s41467-020-20327-5

Cheng, E.H.-Y., Sheiko, T.V., Fisher, J.K., Craigen, W.J., Korsmeyer, S.J., 2003. VDAC2 Inhibits BAK Activation and Mitochondrial Apoptosis. Science 301, 513–517. 10.1126/science.1083995

Clark, D.J., Dhanasekaran, S.M., Petralia, F., Pan, J., Song, X., Hu, Y., da Veiga Leprevost, F., Reva, B., Lih, T.-S.M., Chang, H.-Y., Ma, W., Huang, C., Ricketts, C.J., Chen, L., Krek, A., Li, Y., Rykunov, D., Li, Q.K., Chen, L.S., Ozbek, U., Vasaikar, S., Wu, Y., Yoo, S., Chowdhury, S., Wyczalkowski, M.A., Ji, J., Schnaubelt, M., Kong, A., Sethuraman, S., Avtonomov, D.M., Ao, M., Colaprico, A., Cao, S., Cho, K.-C., Kalayci, S., Ma, S., Liu, W., Ruggles, K., Calinawan, A., Gümüş, Z.H., Geiszler, D., Kawaler, E., Teo, G.C., Wen, B., Zhang, Y., Keegan, S., Li, K., Chen, F., Edwards, N., Pierorazio, P.M., Chen, X.S., Pavlovich, C.P., Hakimi, A.A., Brominski, G., Hsieh, J.J., Antczak, A., Omelchenko, T., Lubinski, J., Wiznerowicz, M., Linehan, W.M., Kinsinger, C.R., Thiagarajan, M., Boja, E.S., Mesri, M., Hiltke, T., Robles, A.I., Rodriguez, H., Qian, J., Fenyö, D., Zhang, B., Ding, L., Schadt, E., Chinnaiyan, A.M., Zhang, Z., Omenn, G.S., Cieslik, M., Chan, D.W., Nesvizhskii, A.I., Wang, P., Zhang, H., Clinical Proteomic Tumor Analysis Consortium, 2019. Integrated Proteogenomic Characterization of Clear Cell Renal Cell Carcinoma. Cell 179, 964–983.e31. 10.1016/j.cell.2019.10.007

Cox, E.J., Marsh, S.A., 2014. A systematic review of fetal genes as biomarkers of cardiac hypertrophy in rodent models of diabetes. PloS One 9, e92903. 10.1371/journal.pone.0092903

Currie, J., Manda, V., Robinson, S.K., Lai, C., Agnihotri, V., Hidalgo, V., Ludwig, R.W., Zhang, K., Pavelka, J., Wang, Z.V., Rhee, J.-W., Lam, M.P.Y., Lau, E., 2024. Simultaneous proteome localization and turnover analysis reveals spatiotemporal features of protein homeostasis disruptions. Nat. Commun. 15, 2207. 10.1038/s41467-024-46600-5

Dai, C., Li, Q., May, H.I., Li, C., Zhang, G., Sharma, G., Sherry, A.D., Malloy, C.R., Khemtong, C., Zhang, Y., Deng, Y., Gillette, T.G., Xu, J., Scadden, D.T., Wang, Z.V., 2020. Lactate Dehydrogenase A Governs Cardiac Hypertrophic Growth in Response to Hemodynamic Stress. Cell Rep. 32, 108087. 10.1016/j.celrep.2020.108087

D’Antonio, M., Nguyen, J.P., Arthur, T.D., Matsui, H., Donovan, M.K.R., D’Antonio-Chronowska, A., Frazer, K.A., 2022. In heart failure reactivation of RNA-binding proteins is associated with the expression of 1,523 fetal-specific isoforms. PLOS Comput. Biol. 18, e1009918. 10.1371/journal.pcbi.1009918

DeLaughter, D.M., Bick, A.G., Wakimoto, H., McKean, D., Gorham, J.M., Kathiriya, I.S., Hinson, J.T., Homsy, J., Gray, J., Pu, W., Bruneau, B.G., Seidman, J.G., Seidman, C.E., 2016. Single-Cell Resolution of Temporal Gene Expression during Heart Development. Dev. Cell 39, 480–490. 10.1016/j.devcel.2016.10.001

Dimasi, C.G., Darby, J.R.T., Morrison, J.L., 2023. A change of heart: understanding the mechanisms regulating cardiac proliferation and metabolism before and after birth. J. Physiol. 601, 1319–1341. 10.1113/JP284137

Dirkx, E., Da Costa Martins, P.A., De Windt, L.J., 2013. Regulation of fetal gene expression in heart failure. Biochim. Biophys. Acta BBA - Mol. Basis Dis. 1832, 2414–2424. 10.1016/j.bbadis.2013.07.023

Dobin, A., Davis, C.A., Schlesinger, F., Drenkow, J., Zaleski, C., Jha, S., Batut, P., Chaisson, M., Gingeras, T.R., 2013. STAR: ultrafast universal RNA-seq aligner. Bioinforma. Oxf. Engl. 29, 15–21. 10.1093/bioinformatics/bts635

Dostal, V., Wood, S.D., Thomas, C.T., Han, Y., Lau, E., Lam, M.P.Y., 2020. Proteomic signatures of acute oxidative stress response to paraquat in the mouse heart. Sci. Rep. 10, 18440. 10.1038/s41598-020-75505-8

Durinck, S., Spellman, P.T., Birney, E., Huber, W., 2009. Mapping identifiers for the integration of genomic datasets with the R/Bioconductor package biomaRt. Nat. Protoc. 4, 1184–1191. 10.1038/nprot.2009.97

Edwards, W., Greco, T.M., Miner, G.E., Barker, N.K., Herring, L., Cohen, S., Cristea, I.M., Conlon, F.L., 2023. Quantitative proteomic profiling identifies global protein network dynamics in murine embryonic heart development. Dev. Cell 58, 1087–1105.e4. 10.1016/j.devcel.2023.04.011

Finn, R.D., Mistry, J., Tate, J., Coggill, P., Heger, A., Pollington, J.E., Gavin, O.L., Gunasekaran, P., Ceric, G., Forslund, K., Holm, L., Sonnhammer, E.L.L., Eddy, S.R., Bateman, A., 2010. The Pfam protein families database. Nucleic Acids Res. 38, D211–D222. 10.1093/nar/gkp985

Franco, D., Lamers, W., Moorman, A.F., 1998. Patterns of expression in the developing myocardium: towards a morphologically integrated transcriptional model. Cardiovasc. Res. 38, 25–53. 10.1016/S0008-6363(97)00321-0

Frankish, A., Diekhans, M., Jungreis, I., Lagarde, J., Loveland, J.E., Mudge, J.M., Sisu, C., Wright, J.C., Armstrong, J., Barnes, I., Berry, A., Bignell, A., Boix, C., Carbonell Sala, S., Cunningham, F., Di Domenico, T., Donaldson, S., Fiddes, I.T., García Girón, C., Gonzalez, J.M., Grego, T., Hardy, M., Hourlier, T., Howe, K.L., Hunt, T., Izuogu, O.G., Johnson, R., Martin, F.J., Martínez, L., Mohanan, S., Muir, P., Navarro, F.C.P., Parker, A., Pei, B., Pozo, F., Riera, F.C., Ruffier, M., Schmitt, B.M., Stapleton, E., Suner, M.-M., Sycheva, I., Uszczynska-Ratajczak, B., Wolf, M.Y., Xu, J., Yang, Y.T., Yates, A., Zerbino, D., Zhang, Y., Choudhary, J.S., Gerstein, M., Guigó, R., Hubbard, T.J.P., Kellis, M., Paten, B., Tress, M.L., Flicek, P., 2021. GENCODE 2021. Nucleic Acids Res. 49, D916–D923. 10.1093/nar/gkaa1087

Fujimura, K., Kano, F., Murata, M., 2008. Identification of PCBP2, a facilitator of IRES-mediated translation, as a novel constituent of stress granules and processing bodies. RNA N. Y. N 14, 425–431. 10.1261/rna.780708

Galdos, F.X., Guo, Y., Paige, S.L., VanDusen, N.J., Wu, S.M., Pu, W.T., 2017. Cardiac Regeneration: Lessons From Development. Circ. Res. 120, 941–959. 10.1161/CIRCRESAHA.116.309040

Gan, P., Wang, Z., Morales, M.G., Zhang, Y., Bassel-Duby, R., Liu, N., Olson, E.N., 2022. RBPMS is an RNA-binding protein that mediates cardiomyocyte binucleation and cardiovascular development. Dev. Cell 57, 959–973.e7. 10.1016/j.devcel.2022.03.017

Geiger, T., Velic, A., Macek, B., Lundberg, E., Kampf, C., Nagaraj, N., Uhlen, M., Cox, J., Mann, M., 2013. Initial quantitative proteomic map of 28 mouse tissues using the SILAC mouse. Mol. Cell. Proteomics MCP 12, 1709–1722. 10.1074/mcp.M112.024919

Gelens, L., Qian, J., Bollen, M., Saurin, A.T., 2018. The Importance of Kinase–Phosphatase Integration: Lessons from Mitosis. Trends Cell Biol. 28, 6–21. 10.1016/j.tcb.2017.09.005

Gillespie, M., Jassal, B., Stephan, R., Milacic, M., Rothfels, K., Senff-Ribeiro, A., Griss, J., Sevilla, C., Matthews, L., Gong, C., Deng, C., Varusai, T., Ragueneau, E., Haider, Y., May, B., Shamovsky, V., Weiser, J., Brunson, T., Sanati, N., Beckman, L., Shao, X., Fabregat, A., Sidiropoulos, K., Murillo, J., Viteri, G., Cook, J., Shorser, S., Bader, G., Demir, E., Sander, C., Haw, R., Wu, G., Stein, L., Hermjakob, H., D’Eustachio, P., 2022. The reactome pathway knowledgebase 2022. Nucleic Acids Res. 50, D687–D692. 10.1093/nar/gkab1028

Gu, Y., Zhou, Y., Ju, S., Liu, X., Zhang, Z., Guo, J., Gao, J., Zang, J., Sun, H., Chen, Q., Wang, J., Xu, J., Xu, Y., Chen, Y., Guo, Y., Dai, J., Ma, H., Wang, C., Jin, G., Li, C., Xia, Y., Shen, H., Yang, Y., Guo, X., Hu, Z., 2022. Multi-omics profiling visualizes dynamics of cardiac development and functions. Cell Rep. 41, 111891. 10.1016/j.celrep.2022.111891

Gygi, S.P., Rochon, Y., Franza, B.R., Aebersold, R., 1999. Correlation between protein and mRNA abundance in yeast. Mol. Cell. Biol. 19, 1720–1730. 10.1128/MCB.19.3.1720

Hammond, D.E., Simpson, D.M., Franco, C., Wright Muelas, M., Waters, J., Ludwig, R.W., Prescott, M.C., Hurst, J.L., Beynon, R.J., Lau, E., 2022. Harmonizing Labeling and Analytical Strategies to Obtain Protein Turnover Rates in Intact Adult Animals. Mol. Cell. Proteomics 21, 100252. 10.1016/j.mcpro.2022.100252

Han, Y., Wennersten, S.A., Wright, J.M., Ludwig, R.W., Lau, E., Lam, M.P.Y., 2022. Proteogenomics reveals sex-biased aging genes and coordinated splicing in cardiac aging. Am. J. Physiol.-Heart Circ. Physiol. 323, H538–H558. 10.1152/ajpheart.00244.2022

Han, Y., Wright, J.M., Lau, E., Lam, M.P.Y., 2020. Determining Alternative Protein Isoform Expression Using RNA Sequencing and Mass Spectrometry. STAR Protoc. 1, 100138. 10.1016/j.xpro.2020.100138

Hart, C.M., Cuvier, O., Laemmli, U.K., 1999. Evidence for an antagonistic relationship between the boundary element-associated factor BEAF and the transcription factor DREF. Chromosoma 108, 375–383. 10.1007/s004120050389

Hasegawa, Y., Struhl, K., 2021. Different SP1 binding dynamics at individual genomic loci in human cells. Proc. Natl. Acad. Sci. 118, e2113579118. 10.1073/pnas.2113579118

Hasman, M., Mayr, M., Theofilatos, K., 2023. Uncovering Protein Networks in Cardiovascular Proteomics. Mol. Cell. Proteomics 22, 100607. 10.1016/j.mcpro.2023.100607

Huang, Q., Szklarczyk, D., Wang, M., Simonovic, M., Von Mering, C., 2023. PaxDb 5.0: Curated Protein Quantification Data Suggests Adaptive Proteome Changes in Yeasts. Mol. Cell. Proteomics 22, 100640. 10.1016/j.mcpro.2023.100640

Hunkeler, N.M., Kullman, J., Murphy, A.M., 1991. Troponin I isoform expression in human heart. Circ. Res. 69, 1409–1414. 10.1161/01.RES.69.5.1409

Huttlin, E.L., Jedrychowski, M.P., Elias, J.E., Goswami, T., Rad, R., Beausoleil, S.A., Villén, J., Haas, W., Sowa, M.E., Gygi, S.P., 2010. A tissue-specific atlas of mouse protein phosphorylation and expression. Cell 143, 1174–1189. 10.1016/j.cell.2010.12.001

Ishii, T., Igawa, T., Hayakawa, H., Fujita, T., Sekiguchi, M., Nakabeppu, Y., 2020. PCBP1 and PCBP2 both bind heavily oxidized RNA but cause opposing outcomes, suppressing or increasing apoptosis under oxidative conditions. J. Biol. Chem. 295, 12247–12261. 10.1074/jbc.RA119.011870

Israelsen, W.J., Vander Heiden, M.G., 2015. Pyruvate kinase: Function, regulation and role in cancer. Semin. Cell Dev. Biol. 43, 43–51. 10.1016/j.semcdb.2015.08.004

Jiang, J., Wu, H., Ji, Y., Han, K., Tang, J.-M., Hu, S., Lei, W., 2024. Development and disease-specific regulation of RNA splicing in cardiovascular system. Front. Cell Dev. Biol. 12, 1423553. 10.3389/fcell.2024.1423553

Karam, J.A.Q., Fréreux, C., Mohanty, B.K., Dalton, A.C., Dincman, T.A., Palanisamy, V., Howley, B.V., Howe, P.H., 2024. The RNA-binding protein PCBP1 modulates transcription by recruiting the G-quadruplex-specific helicase DHX9. J. Biol. Chem. 300, 107830. 10.1016/j.jbc.2024.107830

Karpov, O.A., Stotland, A., Raedschelders, K., Chazarin, B., Ai, L., Murray, C.I., Van Eyk, J.E., 2024. Proteomics of the heart. Physiol. Rev. 104, 931–982. 10.1152/physrev.00026.2023

Keller, A., Rouzeau, J.D., Farhadian, F., Wisnewsky, C., Marotte, F., Lamande, N., Samuel, J.L., Schwartz, K., Lazar, M., Lucas, M., 1995. Differential expression of alpha- and beta-enolase genes during rat heart development and hypertrophy. Am. J. Physiol.-Heart Circ. Physiol. 269, H1843–H1851. 10.1152/ajpheart.1995.269.6.H1843

Klann, K., Tascher, G., Münch, C., 2020. Functional Translatome Proteomics Reveal Converging and Dose-Dependent Regulation by mTORC1 and eIF2α. Mol. Cell 77, 913–925.e4. 10.1016/j.molcel.2019.11.010

Kong, A.T., Leprevost, F.V., Avtonomov, D.M., Mellacheruvu, D., Nesvizhskii, A.I., 2017. MSFragger: ultrafast and comprehensive peptide identification in mass spectrometry-based proteomics. Nat. Methods 14, 513–520. 10.1038/nmeth.4256

Kornienko, J., Rodríguez-Martínez, M., Fenzl, K., Hinze, F., Schraivogel, D., Grosch, M., Tunaj, B., Lindenhofer, D., Schraft, L., Kueblbeck, M., Smith, E., Mao, C., Brown, E., Owens, A., Saguner, A.M., Meder, B., Parikh, V., Gotthardt, M., Steinmetz, L.M., 2023. Mislocalization of pathogenic RBM20 variants in dilated cardiomyopathy is caused by loss-of-interaction with Transportin-3. Nat. Commun. 14, 4312. 10.1038/s41467-023-39965-6

Kory, N., Wyant, G.A., Prakash, G., Uit de Bos, J., Bottanelli, F., Pacold, M.E., Chan, S.H., Lewis, C.A., Wang, T., Keys, H.R., Guo, Y.E., Sabatini, D.M., 2018. SFXN1 is a mitochondrial serine transporter required for one-carbon metabolism. Science 362, eaat9528. 10.1126/science.aat9528

Krikun, G., Schatz, F., Mackman, N., Guller, S., Demopoulos, R., Lockwood, C.J., 2000. Regulation of Tissue Factor Gene Expression In Human Endometrium by Transcription Factors Sp1 and Sp3. Mol. Endocrinol. 14, 393–400. 10.1210/mend.14.3.0430

Kuwahara, K., Nishikimi, T., Nakao, K., 2012. Transcriptional Regulation of the Fetal Cardiac Gene Program. J. Pharmacol. Sci. 119, 198–203. 10.1254/jphs.12R04CP

Lam, M.P.Y., Wang, D., Lau, E., Liem, D.A., Kim, A.K., Ng, D.C.M., Liang, X., Bleakley, B.J., Liu, C., Tabaraki, J.D., Cadeiras, M., Wang, Y., Deng, M.C., Ping, P., 2014. Protein kinetic signatures of the remodeling heart following isoproterenol stimulation. J. Clin. Invest. 124, 1734–1744. 10.1172/JCI73787

Larocque, D., Fragoso, G., Huang, J., Mushynski, W.E., Loignon, M., Richard, S., Almazan, G., 2009. The QKI-6 and QKI-7 RNA Binding Proteins Block Proliferation and Promote Schwann Cell Myelination. PLoS ONE 4, e5867. 10.1371/journal.pone.0005867

Lau, E., Cao, Q., Ng, D.C.M., Bleakley, B.J., Dincer, T.U., Bot, B.M., Wang, D., Liem, D.A., Lam, M.P.Y., Ge, J., Ping, P., 2016. A large dataset of protein dynamics in the mammalian heart proteome. Sci. Data 3, 160015. 10.1038/sdata.2016.15

Lau, E., Han, Y., Williams, D.R., Thomas, C.T., Shrestha, R., Wu, J.C., Lam, M.P.Y., 2019. Splice-Junction-Based Mapping of Alternative Isoforms in the Human Proteome. Cell Rep. 29, 3751–3765.e5. 10.1016/j.celrep.2019.11.026

Lazear, M.R., 2023. Sage: An Open-Source Tool for Fast Proteomics Searching and Quantification at Scale. J. Proteome Res. 22, 3652–3659. 10.1021/acs.jproteome.3c00486

Li, G.-W., Burkhardt, D., Gross, C., Weissman, J.S., 2014. Quantifying absolute protein synthesis rates reveals principles underlying allocation of cellular resources. Cell 157, 624–635. 10.1016/j.cell.2014.02.033

Love, M.I., Huber, W., Anders, S., 2014. Moderated estimation of fold change and dispersion for RNA-seq data with DESeq2. Genome Biol. 15, 550. 10.1186/s13059-014-0550-8

Lowes, B.D., Gilbert, E.M., Abraham, W.T., Minobe, W.A., Larrabee, P., Ferguson, D., Wolfel, E.E., Lindenfeld, J., Tsvetkova, T., Robertson, A.D., Quaife, R.A., Bristow, M.R., 2002. Myocardial Gene Expression in Dilated Cardiomyopathy Treated with Beta-Blocking Agents. N. Engl. J. Med. 346, 1357–1365. 10.1056/NEJMoa012630

Ludwig, R.W., Lau, E., 2021. JCAST: Sample-specific protein isoform databases for mass spectrometry-based proteomics experiments. Softw. Impacts 10, 100163. 10.1016/j.simpa.2021.100163

Lyons, G.E., Schiaffino, S., Sassoon, D., Barton, P., Buckingham, M., 1990. Developmental regulation of myosin gene expression in mouse cardiac muscle. J. Cell Biol. 111, 2427–2436. 10.1083/jcb.111.6.2427

Mantica, F., Iñiguez, L.P., Marquez, Y., Permanyer, J., Torres-Mendez, A., Cruz, J., Franch-Marro, X., Tulenko, F., Burguera, D., Bertrand, S., Doyle, T., Nouzova, M., Currie, P.D., Noriega, F.G., Escriva, H., Arnone, M.I., Albertin, C.B., Wotton, K.R., Almudi, I., Martin, D., Irimia, M., 2024. Evolution of tissue-specific expression of ancestral genes across vertebrates and insects. Nat. Ecol. Evol. 8, 1140–1153. 10.1038/s41559-024-02398-5

Mantica, F., Irimia, M., 2025. Gene Duplication and Alternative Splicing as Evolutionary Drivers of Proteome Specialization. BioEssays e202400202. 10.1002/bies.202400202

Mantica, F., Irimia, M., 2022. The 3D-Evo Space: Evolution of Gene Expression and Alternative Splicing Regulation. Annu. Rev. Genet. 56, 315–337. 10.1146/annurev-genet-071719-020653

Martin, F.J., Amode, M.R., Aneja, A., Austine-Orimoloye, O., Azov, A.G., Barnes, I., Becker, A., Bennett, R., Berry, A., Bhai, J., Bhurji, S.K., Bignell, A., Boddu, S., Branco Lins, P.R., Brooks, L., Ramaraju, S.B., Charkhchi, M., Cockburn, A., Da Rin Fiorretto, L., Davidson, C., Dodiya, K., Donaldson, S., El Houdaigui, B., El Naboulsi, T., Fatima, R., Giron, C.G., Genez, T., Ghattaoraya, G.S., Martinez, J.G., Guijarro, C., Hardy, M., Hollis, Z., Hourlier, T., Hunt, T., Kay, M., Kaykala, V., Le, T., Lemos, D., Marques-Coelho, D., Marugán, J.C., Merino, G.A., Mirabueno, L.P., Mushtaq, A., Hossain, S.N., Ogeh, D.N., Sakthivel, M.P., Parker, A., Perry, M., Piližota, I., Prosovetskaia, I., Pérez-Silva, J.G., Salam, A.I.A., Saraiva-Agostinho, N., Schuilenburg, H., Sheppard, D., Sinha, S., Sipos, B., Stark, W., Steed, E., Sukumaran, R., Sumathipala, D., Suner, M.-M., Surapaneni, L., Sutinen, K., Szpak, M., Tricomi, F.F., Urbina-Gómez, D., Veidenberg, A., Walsh, T.A., Walts, B., Wass, E., Willhoft, N., Allen, J., Alvarez-Jarreta, J., Chakiachvili, M., Flint, B., Giorgetti, S., Haggerty, L., Ilsley, G.R., Loveland, J.E., Moore, B., Mudge, J.M., Tate, J., Thybert, D., Trevanion, S.J., Winterbottom, A., Frankish, A., Hunt, S.E., Ruffier, M., Cunningham, F., Dyer, S., Finn, R.D., Howe, K.L., Harrison, P.W., Yates, A.D., Flicek, P., 2023. Ensembl 2023. Nucleic Acids Res. 51, D933–D941. 10.1093/nar/gkac958

Masamha, C.P., Xia, Z., Peart, N., Collum, S., Li, W., Wagner, E.J., Shyu, A.-B., 2016. CFIm25 regulates glutaminase alternative terminal exon definition to modulate miR-23 function. RNA N. Y. N 22, 830–838. 10.1261/rna.055939.116

McNally, E.M., Kraft, R., Bravo-Zehnder, M., Taylor, D.A., Leinwand, L.A., 1989. Full-length rat alpha and beta cardiac myosin heavy chain sequences. Comparisons suggest a molecular basis for functional differences. J. Mol. Biol. 210, 665–671. 10.1016/0022-2836(89)90141-1

Montañés-Agudo, P., Pinto, Y.M., Creemers, E.E., 2023. Splicing factors in the heart: Uncovering shared and unique targets. J. Mol. Cell. Cardiol. 179, 72–79. 10.1016/j.yjmcc.2023.04.003

Mulligan, G.J., Guo, W., Wormsley, S., Helfman, D.M., 1992. Polypyrimidine tract binding protein interacts with sequences involved in alternative splicing of beta-tropomyosin pre-mRNA. J. Biol. Chem. 267, 25480–25487. 10.1016/S0021-9258(19)74066-6

Oka, T., Xu, J., Molkentin, J.D., 2007. Re-employment of developmental transcription factors in adult heart disease. Semin. Cell Dev. Biol. 18, 117–131. 10.1016/j.semcdb.2006.11.012

Payne, S.H., 2015. The utility of protein and mRNA correlation. Trends Biochem. Sci. 40, 1–3. 10.1016/j.tibs.2014.10.010

Pertea, M., Pertea, G.M., Antonescu, C.M., Chang, T.-C., Mendell, J.T., Salzberg, S.L., 2015. StringTie enables improved reconstruction of a transcriptome from RNA-seq reads. Nat. Biotechnol. 33, 290–295. 10.1038/nbt.3122

Pope, B., Hoh, J.F., Weeds, A., 1980. The ATPase activities of rat cardiac myosin isoenzymes. FEBS Lett. 118, 205–208. 10.1016/0014-5793(80)80219-5

Rajabi, M., Kassiotis, C., Razeghi, P., Taegtmeyer, H., 2007. Return to the fetal gene program protects the stressed heart: a strong hypothesis. Heart Fail. Rev. 12, 331–343. 10.1007/s10741-007-9034-1

Razeghi, P., Young, M.E., Alcorn, J.L., Moravec, C.S., Frazier, O.H., Taegtmeyer, H., 2001. Metabolic Gene Expression in Fetal and Failing Human Heart. Circulation 104, 2923–2931. 10.1161/hc4901.100526

Read, J.A., Winter, V.J., Eszes, C.M., Sessions, R.B., Brady, R.L., 2001. Structural basis for altered activity of M- and H-isozyme forms of human lactate dehydrogenase. Proteins 43, 175–185. 10.1002/1097-0134(20010501)43:2<175::aid-prot1029>3.0.co;2-#

Rees, M.L., Subramaniam, J., Li, Y., Hamilton, D.J., Frazier, O.H., Taegtmeyer, H., 2015. A PKM2 signature in the failing heart. Biochem. Biophys. Res. Commun. 459, 430–436. 10.1016/j.bbrc.2015.02.122

Reiser, P.J., Portman, M.A., Ning, X.-H., Moravec, C.S., 2001. Human cardiac myosin heavy chain isoforms in fetal and failing adult atria and ventricles. Am. J. Physiol.-Heart Circ. Physiol. 280, H1814–H1820. 10.1152/ajpheart.2001.280.4.H1814

Ritchie, M.E., Phipson, B., Wu, D., Hu, Y., Law, C.W., Shi, W., Smyth, G.K., 2015. limma powers differential expression analyses for RNA-sequencing and microarray studies. Nucleic Acids Res. 43, e47. 10.1093/nar/gkv007

Roberts, D.S., Loo, J.A., Tsybin, Y.O., Liu, X., Wu, S., Chamot-Rooke, J., Agar, J.N., Paša-Tolić, L., Smith, L.M., Ge, Y., 2024. Top-down proteomics. Nat. Rev. Methods Primer 4, 38. 10.1038/s43586-024-00318-2

Savitski, M.M., Wilhelm, M., Hahne, H., Kuster, B., Bantscheff, M., 2015. A Scalable Approach for Protein False Discovery Rate Estimation in Large Proteomic Data Sets. Mol. Cell. Proteomics MCP 14, 2394–2404. 10.1074/mcp.M114.046995

Savitski, M.M., Zinn, N., Faelth-Savitski, M., Poeckel, D., Gade, S., Becher, I., Muelbaier, M., Wagner, A.J., Strohmer, K., Werner, T., Melchert, S., Petretich, M., Rutkowska, A., Vappiani, J., Franken, H., Steidel, M., Sweetman, G.M., Gilan, O., Lam, E.Y.N., Dawson, M.A., Prinjha, R.K., Grandi, P., Bergamini, G., Bantscheff, M., 2018. Multiplexed Proteome Dynamics Profiling Reveals Mechanisms Controlling Protein Homeostasis. Cell 173, 260–274.e25. 10.1016/j.cell.2018.02.030

Schwanhäusser, B., Busse, D., Li, N., Dittmar, G., Schuchhardt, J., Wolf, J., Chen, W., Selbach, M., 2011. Global quantification of mammalian gene expression control. Nature 473, 337–342. 10.1038/nature10098

Searle, B.C., Yergey, A.L., 2020. An efficient solution for resolving iTRAQ and TMT channel cross-talk. J. Mass Spectrom. 55, e4354. 10.1002/jms.4354

Smink, J.J., Bégay, V., Schoenmaker, T., Sterneck, E., De Vries, T.J., Leutz, A., 2009. Transcription factor C/EBPβ isoform ratio regulates osteoclastogenesis through MafB. EMBO J. 28, 1769– 1781. 10.1038/emboj.2009.127

Srivastava, H., Lippincott, M.J., Currie, J., Canfield, R., Lam, M.P.Y., Lau, E., 2022. Protein prediction models support widespread post-transcriptional regulation of protein abundance by interacting partners. PLOS Comput. Biol. 18, e1010702. 10.1371/journal.pcbi.1010702

Suhre, K., 2024. Genetic associations with ratios between protein levels detect new pQTLs and reveal protein-protein interactions. Cell Genomics 4, 100506. 10.1016/j.xgen.2024.100506

Szklarczyk, D., Kirsch, R., Koutrouli, M., Nastou, K., Mehryary, F., Hachilif, R., Gable, A.L., Fang, T., Doncheva, N.T., Pyysalo, S., Bork, P., Jensen, L.J., von Mering, C., 2023. The STRING database in 2023: protein–protein association networks and functional enrichment analyses for any sequenced genome of interest. Nucleic Acids Res. 51, D638–D646. 10.1093/nar/gkac1000

Taegtmeyer, H., Sen, S., Vela, D., 2010. Return to the fetal gene program: a suggested metabolic link to gene expression in the heart. Ann. N. Y. Acad. Sci. 1188, 191–198. 10.1111/j.1749-6632.2009.05100.x

Talman, V., Teppo, J., Pöhö, P., Movahedi, P., Vaikkinen, A., Karhu, S.T., Trošt, K., Suvitaival, T., Heikkonen, J., Pahikkala, T., Kotiaho, T., Kostiainen, R., Varjosalo, M., Ruskoaho, H., 2018. Molecular Atlas of Postnatal Mouse Heart Development. J. Am. Heart Assoc. 7, e010378. 10.1161/JAHA.118.010378

The, M., MacCoss, M.J., Noble, W.S., Käll, L., 2016. Fast and Accurate Protein False Discovery Rates on Large-Scale Proteomics Data Sets with Percolator 3.0. J. Am. Soc. Mass Spectrom. 27, 1719– 1727. 10.1007/s13361-016-1460-7

The UniProt Consortium, Bateman, A., Martin, M.-J., Orchard, S., Magrane, M., Ahmad, S., Alpi, E., Bowler-Barnett, E.H., Britto, R., Bye-A-Jee, H., Cukura, A., Denny, P., Dogan, T., Ebenezer, T., Fan, J., Garmiri, P., Da Costa Gonzales, L.J., Hatton-Ellis, E., Hussein, A., Ignatchenko, A., Insana, G., Ishtiaq, R., Joshi, V., Jyothi, D., Kandasaamy, S., Lock, A., Luciani, A., Lugaric, M., Luo, J., Lussi, Y., MacDougall, A., Madeira, F., Mahmoudy, M., Mishra, A., Moulang, K., Nightingale, A., Pundir, S., Qi, G., Raj, S., Raposo, P., Rice, D.L., Saidi, R., Santos, R., Speretta, E., Stephenson, J., Totoo, P., Turner, E., Tyagi, N., Vasudev, P., Warner, K., Watkins, X., Zaru, R., Zellner, H., Bridge, A.J., Aimo, L., Argoud-Puy, G., Auchincloss, A.H., Axelsen, K.B., Bansal, P., Baratin, D., Batista Neto, T.M., Blatter, M.-C., Bolleman, J.T., Boutet, E., Breuza, L., Gil, B.C., Casals-Casas, C., Echioukh, K.C., Coudert, E., Cuche, B., De Castro, E., Estreicher, A., Famiglietti, M.L., Feuermann, M., Gasteiger, E., Gaudet, P., Gehant, S., Gerritsen, V., Gos, A., Gruaz, N., Hulo, C., Hyka-Nouspikel, N., Jungo, F., Kerhornou, A., Le Mercier, P., Lieberherr, D., Masson, P., Morgat, A., Muthukrishnan, V., Paesano, S., Pedruzzi, I., Pilbout, S., Pourcel, L., Poux, S., Pozzato, M., Pruess, M., Redaschi, N., Rivoire, C., Sigrist, C.J.A., Sonesson, K., Sundaram, S., Wu, C.H., Arighi, C.N., Arminski, L., Chen, C., Chen, Y., Huang, H., Laiho, K., McGarvey, P., Natale, D.A., Ross, K., Vinayaka, C.R., Wang, Q., Wang, Y., Zhang, J., 2023. UniProt: the Universal Protein Knowledgebase in 2023. Nucleic Acids Res. 51, D523–D531. 10.1093/nar/gkac1052

Thrasher, J.R., Cooper, M.D., Dunaway, G.A., 1981. Developmental changes in heart and muscle phosphofructokinase isozymes. J. Biol. Chem. 256, 7844–7848. 10.1016/S0021-9258(18)43355-8

Vigil-Garcia, M., Demkes, C.J., Eding, J.E.C., Versteeg, D., De Ruiter, H., Perini, I., Kooijman, L., Gladka, M.M., Asselbergs, F.W., Vink, A., Harakalova, M., Bossu, A., Van Veen, T.A.B., Boogerd, C.J., Van Rooij, E., 2021. Gene expression profiling of hypertrophic cardiomyocytes identifies new players in pathological remodelling. Cardiovasc. Res. 117, 1532–1545. 10.1093/cvr/cvaa233

Wang, H., Chen, Y., Li, X., Chen, Guojun, Zhong, L., Chen, Gangbing, Liao, Y., Liao, W., Bin, J., 2016. Genome-wide analysis of alternative splicing during human heart development. Sci. Rep. 6, 35520. 10.1038/srep35520

Wang, H., Dai, C., Pfeuffer, J., Sachsenberg, T., Sanchez, A., Bai, M., Perez-Riverol, Y., 2023. Tissue-based absolute quantification using large-scale TMT and LFQ experiments. PROTEOMICS 23, 2300188. 10.1002/pmic.202300188

Wang, J., Yu, W., D’Anna, R., Przybyla, A., Wilson, M., Sung, M., Bullen, J., Hurt, E., D’Angelo, G., Sidders, B., Lai, Z., Zhong, W., 2023. Pan-Cancer Proteomics Analysis to Identify Tumor-Enriched and Highly Expressed Cell Surface Antigens as Potential Targets for Cancer Therapeutics. Mol. Cell. Proteomics 22, 100626. 10.1016/j.mcpro.2023.100626

Wang, X., Codreanu, S.G., Wen, B., Li, K., Chambers, M.C., Liebler, D.C., Zhang, B., 2018. Detection of Proteome Diversity Resulted from Alternative Splicing is Limited by Trypsin Cleavage Specificity. Mol. Cell. Proteomics 17, 422–430. 10.1074/mcp.RA117.000155

Warren, C.F.A., Wong-Brown, M.W., Bowden, N.A., 2019. BCL-2 family isoforms in apoptosis and cancer. Cell Death Dis. 10, 177. 10.1038/s41419-019-1407-6

White, E.J.F., Matsangos, A.E., Wilson, G.M., 2017. AUF1 regulation of coding and noncoding RNA. WIREs RNA 8, e1393. 10.1002/wrna.1393

Wojtkiewicz, M., Berg Luecke, L., Castro, C., Burkovetskaya, M., Mesidor, R., Gundry, R.L., 2022. Bottom-up proteomic analysis of human adult cardiac tissue and isolated cardiomyocytes. J. Mol. Cell. Cardiol. 162, 20–31. 10.1016/j.yjmcc.2021.08.008

Yang, K.L., Yu, F., Teo, G.C., Li, K., Demichev, V., Ralser, M., Nesvizhskii, A.I., 2023. MSBooster: improving peptide identification rates using deep learning-based features. Nat. Commun. 14, 4539. 10.1038/s41467-023-40129-9

Yang, M., Petralia, F., Li, Z., Li, H., Ma, W., Song, X., Kim, S., Lee, H., Yu, H., Lee, B., Bae, S., Heo, E., Kaczmarczyk, J., Stępniak, P., Warchoł, M., Yu, T., Calinawan, A.P., Boutros, P.C., Payne, S.H., Reva, B., NCI-CPTAC-DREAM Consortium, Boja, E., Rodriguez, H., Stolovitzky, G., Guan, Y., Kang, J., Wang, P., Fenyö, D., Saez-Rodriguez, J., 2020. Community Assessment of the Predictability of Cancer Protein and Phosphoprotein Levels from Genomics and Transcriptomics. Cell Syst. 11, 186–195.e9. 10.1016/j.cels.2020.06.013

Ye, J., Llorian, M., Cardona, M., Rongvaux, A., Moubarak, R.S., Comella, J.X., Bassel-Duby, R., Flavell, R.A., Olson, E.N., Smith, C.W.J., Sanchis, D., 2013. A pathway involving HDAC5, cFLIP and caspases regulates expression of the splicing regulator polypyrimidine tract binding protein in the heart. J. Cell Sci. 126, 1682–1691. 10.1242/jcs.121384

Yoshikawa, S., Nagao, M., Toh, R., Shinohara, M., Iino, T., Irino, Y., Nishimori, M., Tanaka, H., Satomi-Kobayashi, S., Ishida, T., Hirata, K.-I., 2022. Inhibition of glutaminase 1-mediated glutaminolysis improves pathological cardiac remodeling. Am. J. Physiol.-Heart Circ. Physiol. 322, H749–H761. 10.1152/ajpheart.00692.2021

Yu, G., He, Q.-Y., 2016. ReactomePA: an R/Bioconductor package for reactome pathway analysis and visualization. Mol. Biosyst. 12, 477–479. 10.1039/C5MB00663E

Zannoni, G.F., Monterossi, G., De Stefano, I., Gargini, A., Salerno, M.G., Farulla, I., Travaglia, D., Vellone, V.G., Scambia, G., Gallo, D., 2013. The expression ratios of estrogen receptor α (ERα) to estrogen receptor β1 (ERβ1) and ERα to ERβ2 identify poor clinical outcome in endometrioid endometrial cancer. Hum. Pathol. 44, 1047–1054. 10.1016/j.humpath.2012.09.007

Zecha, J., Satpathy, S., Kanashova, T., Avanessian, S.C., Kane, M.H., Clauser, K.R., Mertins, P., Carr, S.A., Kuster, B., 2019. TMT Labeling for the Masses: A Robust and Cost-efficient, In-solution Labeling Approach. Mol. Cell. Proteomics MCP 18, 1468–1478. 10.1074/mcp.TIR119.001385

Zhao, Q., Sun, Q., Zhou, L., Liu, K., Jiao, K., 2019. Complex Regulation of Mitochondrial Function During Cardiac Development. J. Am. Heart Assoc. 8, e012731. 10.1161/JAHA.119.012731

Zhou, T., Pan, J., Xu, K., Yan, C., Yuan, J., Song, H., Han, Y., 2024. Single-cell transcriptomics in MI identify Slc25a4 as a new modulator of mitochondrial malfunction and apoptosis-associated cardiomyocyte subcluster. Sci. Rep. 14, 9274. 10.1038/s41598-024-59975-8

Zhu, A., Ibrahim, J.G., Love, M.I., 2019. Heavy-tailed prior distributions for sequence count data: removing the noise and preserving large differences. Bioinforma. Oxf. Engl. 35, 2084–2092. 10.1093/bioinformatics/bty895

